# Diffusion MRI Tractography Predicts Electrophysiological Connectivity and Explains Spectral Signatures of Evoked Potentials in the Human Brain

**DOI:** 10.64898/2026.09.15.751678

**Authors:** S Shailja, Dian Lyu, Gustavo Chau Loo Kung, Leili Mortazavi, Erpeng Dai, Michael M. Zeineh, Vivek P. Buch, Karl Deisseroth, Josef Parvizi, Jennifer A. McNab

## Abstract

White matter fiber bundles are the structural conduits of information flow in the human brain and thereby mediate the spatial trajectories and timings of the electrophysiological signaling. Here, we combined diffusion magnetic resonance imaging (dMRI) and stereoelectroencephalography (SEEG) recordings in neurosurgical patients to develop an integrated framework for predicting and interpreting causal electrophysiological connectivity patterns between pairs of brain regions. We used repeated single-pulse electrical stimulation in 40 participants implanted with a total of 5794 intracranial electrodes throughout the human brain, encompassing both cortical and multiple thalamic nuclei. A nonlinear, time-frequency manifold learning approach was used to define electrophysiological connectivity, which was then compared with subject-specific and atlas-based structural connectivity. Across 150,000 electrode pairs, we found that the presence of a structural connection predicted causal electrophysiological connectivity with a probability of ≈ 0.95; and its absence predicted the lack of the direct electrophysiological connectivity with a probability of ≈ 0.8. We also show evidence of indirect/polysynaptic pathways supported by both modalities and reported neural features from time-frequency decomposition that distinguished between direct and indirect signaling. We demonstrated that an early phase-locked broadband component (10–70 ms) marked direct structural pathways, whereas delayed and slower components reflected indirect propagation. Notably, we reported that thalamic involvement within an indirect pathway results in increased latency (*>* 200 ms) and enhanced late oscillatory behavior, despite increased conduction velocity measures along thalamo-cortical pathways. Therefore, our multimodal framework maps human brain connectivity, bridging structural architecture, causal electrophysiological dynamics, and network-level communication.

## 1 Introduction

Understanding of the structure-function relationships in the human brain is a fundamental goal of neuroscience [1–4]. Diffusion MRI (dMRI) is the only tool to study the structural connections *in vivo* [5]. Prior work linking tractography with resting-state fMRI [6] and/or surface EEG [7] have been challenging to interpret. Inherently, resting-state fMRI and scalp-EEG measurements are limited in their sensitivity making correlates with structure challenging. Brain function depends on electrophysiological interactions throughout its cortical and subcortical regions [8], which can be measured directly by stereo-electroencephalography (SEEG [9]), an approach used routinely for intracranial monitoring of patients who are candidates for neurosurgical procedures related to their medically refractory epilepsy [10]. Implanted electrodes, often in the range of ≈ 150 contacts, usually cover diverse brain areas to localize seizure onset zones and guide possible surgical interventions [11]. The increased safety afforded by the advent of robotized SEEG procedures [12, 13] has led to increased and broader clinical use. This rich data gathered with SEEG represents a valuable new opportunity to investigate causal relationships among focal brain areas with both high temporal resolution and anatomical precision in the human brain.

Another advantage of SEEG method is that it provides a platform for experiments with high causal relevance [14]. One commonly used causal method is the study of “effective electrophysiological connectivity” in which single electrical pulses are applied repeatedly (≈ 50 times) to a pair of electrode contacts while recording evoked potentials in every other available electrode site, a measure traditionally known as “cortico-cortical evoked potential” (CCEP) [15]. The profile of CCEPs generated in the brain has been used to deduce information about large-scale functional networks of the human brain [16–18], highlighting how CCEP mapping can provide insight into the neural basis of non-invasive imaging signals.

Despite its strengths, SEEG suffers from sparse sampling [19]. By contrast, tractography provides a confidence map of whole brain structural connectivity non-invasively. It is well-established that, within white matter, the orientation with the most rapid diffusive displacement reliably corresponds to the orientation of white matter fiber bundles. Tractography, however, suffers from false positives/negatives, limited sensitivity to small fiber bundles, ambiguity between neighbouring tracts, and challenges tracking in gray matter [20, 21]. To better understand these limitations, prior studies have compared tractography with tract tracer data in animals [22, 23]. We hypothesize that diffusion tractography provides sufficient accuracy to predict and interpret stimulation-evoked causal electrophysiological connectivity in the brain; see Figure 1. Our hypotheses are motivated by the evidence available in the extant literature. For example, Silverstein et. al. [24] developed a dynamic tractography framework that demonstrated that the strength and timing of the “N1” component of CCEPs (10-50 ms post-stimulus) for electrocorticography are dependent on the length and fractional anisotropy of white matter pathways. In a separate study, the CCEP cortical network was compared with resting-state functional and structural connectivity, with distant interactions aligning most closely with tractography-derived structural pathways [25]. It has also been shown that probabilistic mapping of structural connectivity and CCEP features has characterized the dynamics of large fiber bundles [26]. More recently, Śeguin et al. [27] showed that network communication models derived from diffusion MRI structural connectivity may explain multi-step propagations between unconnected brain regions. However, these previous studies have focused predominantly on cortical connectivity, leaving the contribution of the thalamus to brain-wide communication comparatively unexplored. Also, they have largely relied on univariate peak or time-to-peak metrics within fixed windows, which miss the complex dynamics of stimulation evoked responses to distinguish between direct versus indirect connectivity.

**Fig. 1:**
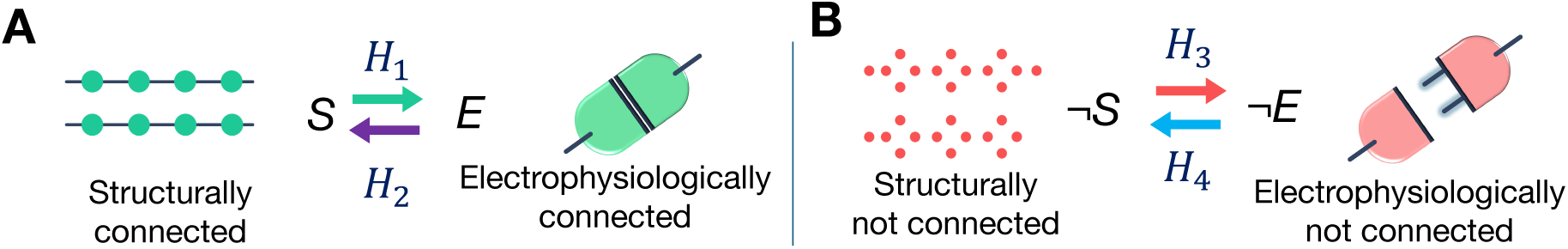
Hypotheses for correlating structural connectivity and causal electrophysiological connectivity. *S* represents structurally connected and ¬*S* denotes structurally “not” connected events, while *E* and ¬*E* denote electrophysiologically connected and not connected events, respectively. The ¬ symbol is the standard logical negation operator denoting “not”. We hypothesize that: **(A)** *H*_1_ & *H*_2_: If a pair of electrode sites are structurally connected, then there is an electrophysiological connection between them, and vice versa. **(B)** *H*_3_ & *H*_4_: If a pair of electrode sites are structurally not connected, then no electrophysiological connection will be observed, and vice versa.

A particularly novel aspect of our study is the inclusion of unprecedented recordings and stimulations from the human thalamus, together with three methodological innovations that distinguish our work from prior studies. Firstly, we define causal electrophysiological connectivity by leveraging the recent developments in the analysis of CCEP by integrating both spectral power and coherence across repeated trials of stimulation [16] instead of relying on arbitrary thresholds to differentiate significant connections from noise. We also expand the temporal window and study spectral changes unfolding in time after the first evoked responses are generated. Secondly, we define structural connectivity in two complimentary ways: (a) patient-specific probabilistic tractography in 10 SEEG subjects, and (b) atlas-based probabilistic tractography using Human Connectome Project (HCP) data (N=1065) [29] for all 40 SEEG subjects. Our approach yielded new information that is currently not available in the literature. Specifically, we demonstrate correspondence between tractography-defined structural connectivity and spectral features of CCEPs including not only cortical connections but also thalamic connections in the human brain. We provide structural pathway information about “direct” versus “indirect” electrophysiological signaling in human brain based on the merging of spectral data from the observed evoked potentials with structural data harnessed from dMRI. Thirdly, we correlate diffusion microstructure features with conduction velocity based on CCEP latency along the tract. Based on our results, we anticipate that non-invasive, whole-brain, *in vivo* prediction of conduction velocity will be possible. Notably, we report that thalamic involvement results in increased latency, despite increased conduction velocity measures along thalamo-cortical pathways. In summary, our work provides a non-invasive framework for testing fundamental hypotheses of structure-function organizations in the human brain. Beyond the clinical application in neurosurgical planning, this work advances our understanding of thalamo-cortical dynamics.

## 2 Results

### 2.1 Multi-model SEEG and diffusion MRI tractography data formalize structural connectivity and causal electrophysiological connectivity

Our results reflect a retrospective study of 40 patients with medically refractory epilepsy (40% female; mean age ± SD: 37.9 ± 12.3) who underwent a SEEG procedure, out of which we acquired dMRI scans for 10 patients prior to their SEEG (Figure 2, Supplementary Table A1). For these subjects we found 143±33 electrodes per subject with a total of 5794 electrode sites across all subjects. Figure 3(B,E) depict the diverse locations of electrodes and the intersections of electrodes with 64 different white matter tracts that have been identified in a tractography atlas and correspond with known anatomy (see Figure 3E). It is this broad electrode coverage across the brain that enables our study of correlating white matter architecture to causal electrophysiological function.

**Fig. 2:**
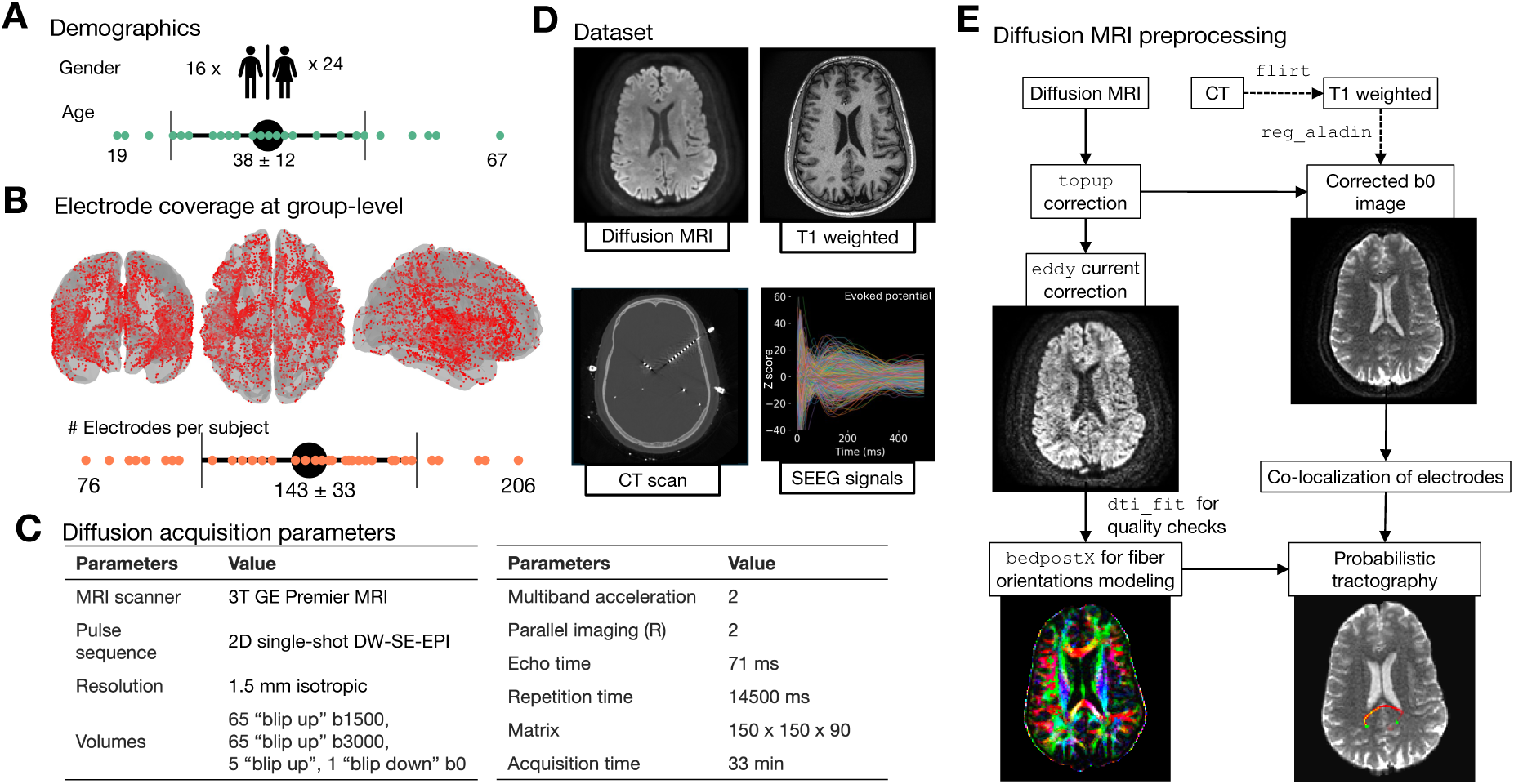
SEEG and diffusion MRI data for electrophysiological and structural information. **(A)** Demographics of 40 medically refractory epilepsy patients who underwent SEEG at the Stanford Hospital. **(B)** The entire dataset encompasses 5794 sites across all subjects, with each adjacent pair stimulated approximately ∼45 times. The centroids of the bipolar pairs are plotted on the FS LR (FreeSurfer Left-Right) surface space for visualization purpose using Nilearn [28]. **(C)** Multi-shell diffusion data with 1.5 mm isotropic resolution were acquired prior to each patient’s SEEG procedure. **(D)** Example imaging and the trial-averaged evoked potentials showing complex and variable electrophysiological responses (local field potential) evoked by the stimulation of a given site. Out of 40 patients, 10 participants were recruited to undergo a separate research diffusion MRI scan prior to their scheduled SEEG procedure. Pre-implant T1 along with the post-implant CT scan and cerebro-cerebral evoked potentials were utilized for 40 patients in this study. **(E)** Diffusion data was preprocessed using FSL. Clinically acquired post-implant CT scans were co-registered with dMRI to determine electrode locations which were used for probabilistic tracking.

**Fig. 3:**
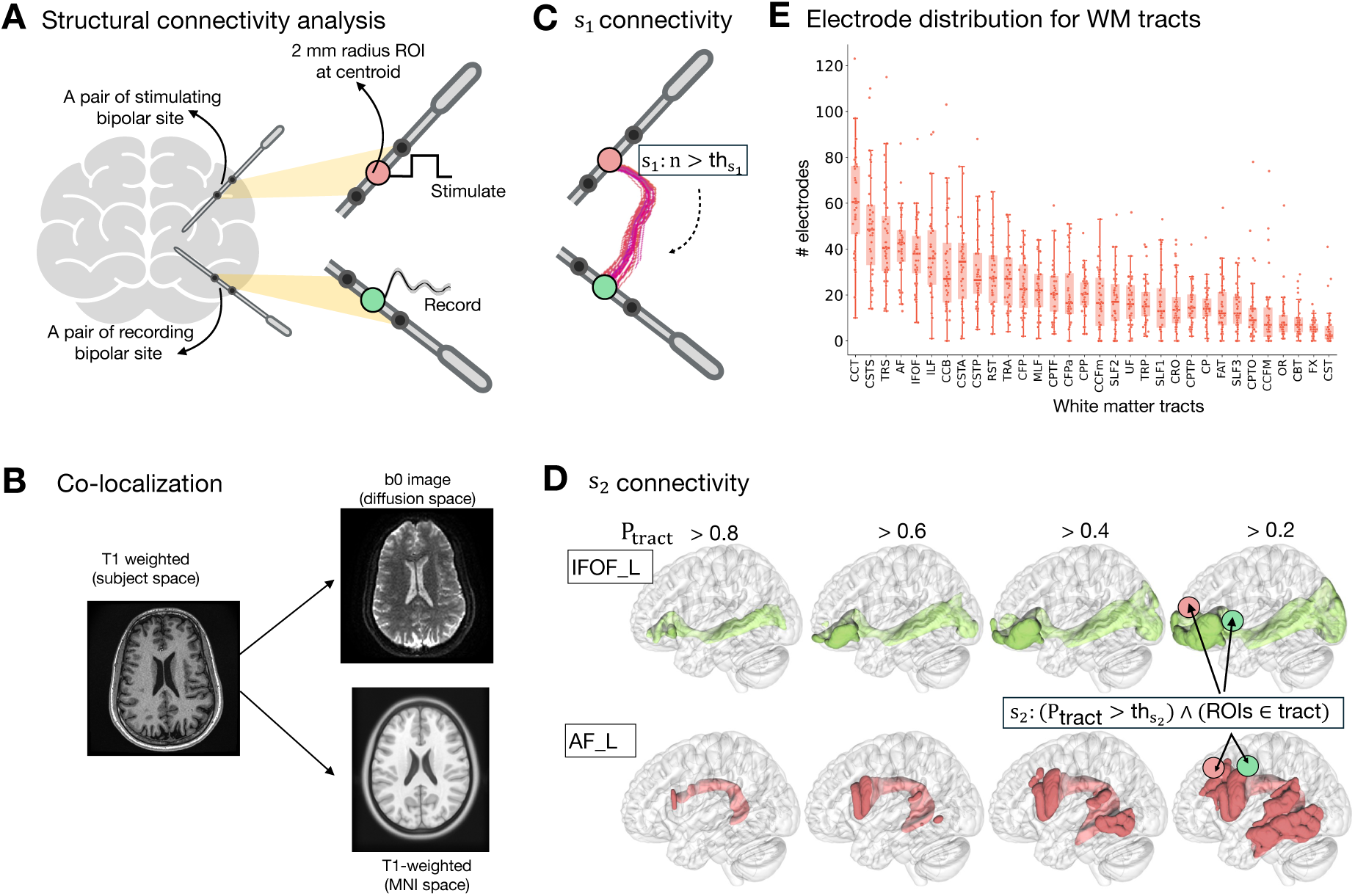
Formalizing the definitions of structural connectivity using patient-specific diffusion MRI and HCP atlas. **(A)** The ROIs of the stimulated and recorded electrodes are 2mm spheres around the centroids of their respective bipolar electrode channels. A pair of stimulating and recording electrodes is *s*_1_-connected if the total streamline count between them is greater than a given streamline threshold (th_s1_). **(B)** Electrode co-localization for **(C)** *s*_1_ connectivity in the patient’s diffusion space and **(D)** *s*_2_ connectivity in MNI space. A pair of stimulating and recording electrodes is *s*_2_-connected if both were located in the same white matter tract at a given tract probability (th_s2_). **(E)** Patient wise distribution of electrodes in white matter tracts from population-probability HCP tractography atlas non-zero overlap and tract probability (th_s2_). Refer to https://brain.labsolver.org/hcp_trk_atlas.html for abbreviations of white matter tracts.

A primary result of our work is a formalism to compare multi-modal connectivity measurements. At the crux of this comparison is establishing biologically meaningful, quantifiable and reproducible definitions of connectivity. We posit two definitions of structural connectivity: 1) *s*_1_-connectivity (N=10) based on the patient-specific diffusion MRI probabilistic tractography (Figure 3A,C) and 2) *s*_2_-connectivity (N=40) based on the intersection of depth electrodes with tracts defined by a probabilistic HCP tractography atlas (Figure 3D). We measured the cerebro-cerebral evoked potentials (CCEP) to define causal electrophysiological connectivity between a pair of electrode sites. To quantify causal electrophysiological connectivity we leveraged a nonlinear manifold learning model trained on the spectral information of evoked potentials across time and frequency domains [16] to classify a pair of electrodes as “connected” vs. “not-connected”. Along with the binary classification of a given pair as electrophysiologically connected or not, this algorithm also identified the significant time/frequency boundaries of three distinct neural features (F1, F2, F3) in the electrophysiology data: (1) F1 in the [30, 70] Hz (gamma) range within 10-70 ms, (2) F2 in the [5, 8] Hz (theta) to [8, 15] Hz (alpha) range, 70-200 ms post-stimulation, and (3) F3 in the theta band with oscillations 200-400 ms post-stimulation.

Our results are based on four fundamental hypotheses:

- *H*_1_: If a structural connection (*S*) is observed, then an electrophysiological connection (*E*) will be observed and vice versa (*H*_2_).
- *H*_3_: If no structural connection (¬*S*) is observed, then no electrophysiological connection (¬*E*) will be observed and vice-versa (*H*_4_).

Following a Bayesian approach [30, 31], we compute *P* (*S* → *E*), which denotes the probability of implication. This is formalized as *P* (*E*|*S*) and the rest follows similarly.

### 2.2 Structural connectivity implies causal electrophysiological connectivity

For *s*_1_-connectivity, based on probabilistic tractography between each electrode pair (2 mm ROIs) with a streamline threshold of th_s_1 (Figure 3A), higher values of th_s_1 imply strong confidence in structural connectivity. The results of our hypothesis testing with varying th_s_1 are shown in Figure 4A. At th_s_1 = 10, across 32, 677 electrode pairs at the group-level (*N* = 10), we observe *S* ∧ *E* = 3289 and *S* ∧ ¬*E* = 172.

**Fig. 4:**
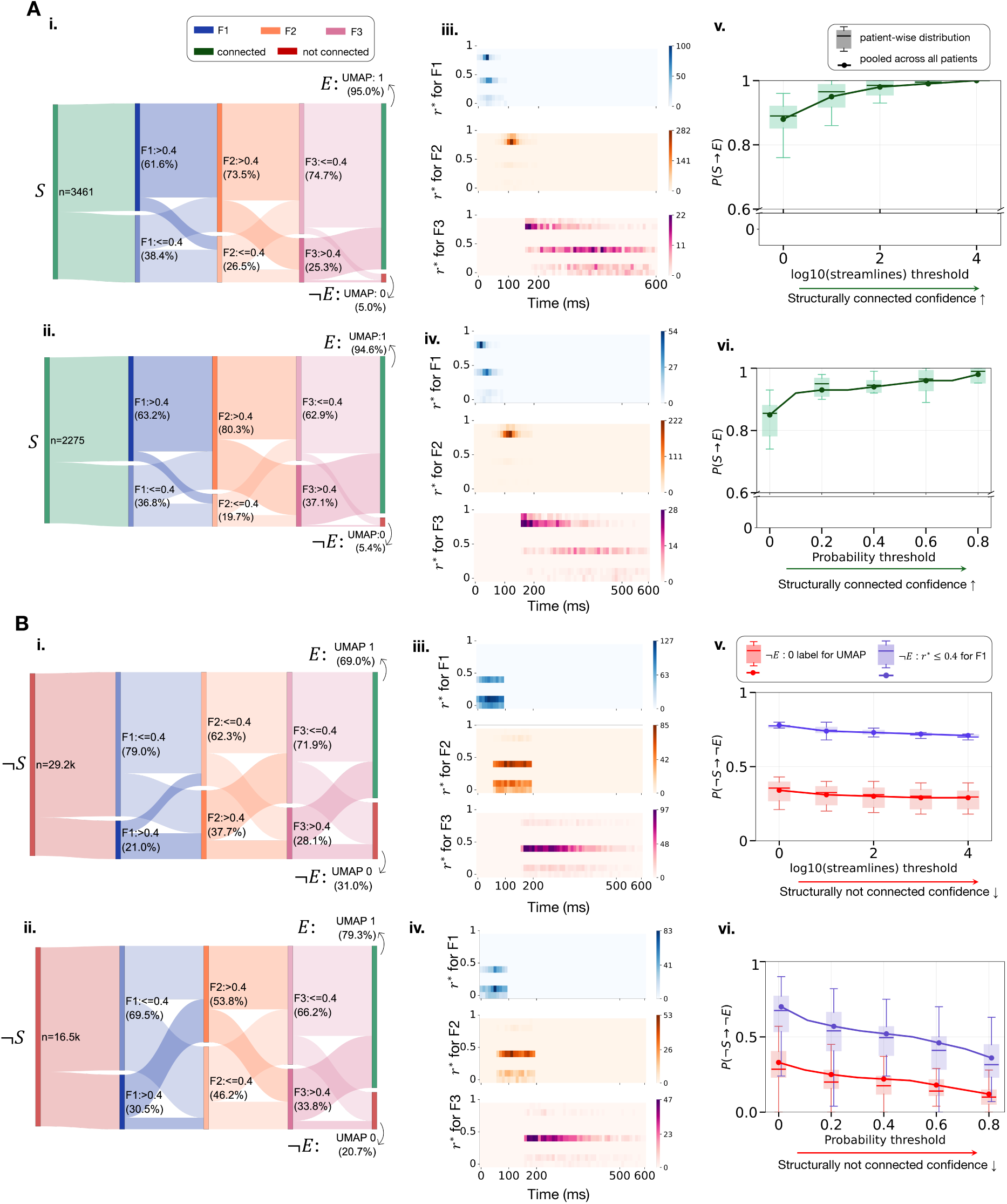
Diffusion MRI tractography predicts causal electrophysiological connectivity. **(A)** Structural connectivity implies causal electrophysiological connectivity. **(i,ii)** Alluvial plots for *s*_1_-connected electrode pairs at *th*_s_1__ = 10 and *s*_2_-connected electrode pairs at *th*_s_2 = 0.5 showing spectral features of the evoked potentials (F1, F2, F3 with threshold on the maximum Pearson correlation coefficient *r*^∗^ = 0.4) and the resulting electro-physiological connectivity label (UMAP:1 = *E*, UMAP:0 = ¬*E*).**(iii,iv)** Time to *r*^∗^ (latency) heatmaps for F1, F2, F3 in *s*_1_ and *s*_2_-connected electrode pairs which are also *E* highlights rapid F1. **(v)** Probability of implication *P* (*E* | *S*) versus log_10_ streamline threshold th_s1_. Increasing the confidence of structural connectivity by increasing the strictness of tractography raises *P* (*E* | *S*) toward 1. **(vi)** *P* (*E* | *S*) versus white matter probability threshold th_s2_ shows the same trend. Solid lines show probability estimates pooled across all subjects while box plots indicate variability of intra-subject probability estimates. **(B)** Structurally not connected electrode pairs lack “direct” electrophysiological connectivity **(i,ii)** Alluvial plots for *s*_1_-not-connected pairs and *s*_2_-not-connected pairs showing spectral features of the evoked potentials (F1, F2, F3; threshold *r*^∗^ = 0.4) and the resulting electrophysiology label (UMAP:1 = *E*, UMAP:0 = ¬*E*). Compared with the connected sets, F1 feature is less prevalent. **(iii,iv)** Time to *r*^∗^ (latency) heatmaps for F1, F2, F3 among not-connected pairs that are correctly labeled ¬*E*. Here, *r*^∗^ is below 0.5 for F1, F2, and F3 with time latency broadly dispersed, unlike the concentrated patterns of F1 and F2 in connected pairs. **(v)** Probability of implication *P* (¬*E* | ¬*S*) versus log_10_ streamline threshold th_s1_ for direct and indirect *E*. Increasing th_s1_ relaxes the definition structurally not connected electrode pairs and thus lower *P* (¬*E* | ¬*S*). **(vi)** *P* (¬*E* | ¬*S*) versus white matter probability threshold th_s2_ shows the same trend. Lines show pooled estimates; box plots indicate variability across subjects. Solid red line plot in **(e)** and **(f)** denotes the *P* (¬*E* | ¬*S*) after removing the possible indirectly connected electrode pairs (*r*∗ *>* 0.4 for F2 or F3).

The probability of *S* implying is *E*: *P* (*E* | *S*) = 0.95 (95% Wilson CI [0.943, 0.957]), indicating that structurally connected electrode pairs were highly likely to exhibit causal electrophysiological connectivity. This association was highly significant (Pearson *χ*^2^= 1033.77, *p <* 0.001), with a risk ratio of RR = 1.4 and an odds ratio of OR = 8.6 and above the baseline (*P* (*E*) = 0.7).

For *s*_2_-connectivity, based on the intersection of electrode ROIs with HCP-atlas defined tract segmentation and voxel-wise probabilities exceeding th_s_2 [29], higher values of th_s_2 imply stronger confidence in structural connectivity. Across 156,061 electrode pairs at th_s_2 = 0.5, we observed *S* ∧ *E* = 2,153 and *S* ∧ ¬*E* = 122, resulting in *P* (*E* | *S*) = 0.946 (95% Wilson CI [0.936, 0.955]). Thus, for *s*_2_ connectivity, similar to *s*_1_, we find that presence of HCP atlas-based structural connectivity between a pair of electrodes implies causal electrophysiological connectivity with ∼ 95% confidence. This association is also highly significant (Pearson *χ*^2^ = 519.45, *p <* 0.001) with RR = 1.29 and OR = 6.37. We also identified electrode pairs that satisfied both the *s*_1_- and *s*_2_-connected criteria. Of the 3,078 such pairs, 2,960 (96%) were electrophysiologically connected, suggesting that agreement between probabilistic tractography and HCP atlas-based tract overlap provides stronger evidence of structural connectivity.

These results strongly support the *H*_1_ hypothesis and indicate that the presence of structural connectivity substantially increases the probability of observing causal electrophysiological connectivity (Fig. 4). Spectral feature distributions for structurally connected pairs show prominent presence of the F1 feature — sharp wave (increased power in high gamma) with tight phase locking to the stimulation onset within the early stage after stimulation within 10 ms to 70 ms post-stimulation. Structurally connected pairs typically have the maximum Pearson correlation coefficient (*r*^∗^) *>* 0.4 for F1 in more than 60% of such pairs (see Figure 4A-i). We also observe that for electrode pairs that are structurally as well as electrophysiologically connected have prominent presence of F1 feature in the early stage (*<* 70 ms) after stimulation. On redefining *S* to be presence of F1 (*r*^∗^ *>* 0.4), *RR* increases to 2.5.

Varying th_s_1 and th_s_2, we observe the trade off between electrode coverage and confidence of structural connectivity (Figure 4A:v,vi). That is, as th_s_1 and th_s_2 increase, fewer pairs are *S* but *P* (*E* | *S*) approaches 1. This is expected as increasing these thresholds provides stricter definitions of structural connectivity. For example, all *s*_1_-connected pairs are electrophysiologically connected at th_s_1 = 10,000. *H*_4_ is the contrapositive of *H*_1_, so we observe similar results for *H*_4_ formalized as *P* (¬*E* → ¬*S*) = *P* (¬*S* | ¬*E*) which is 0.98 for *s*_1_-connectivity and 0.97 for *s*_2_-connectivity. The results for *H*_4_ show that whenever electrode sites in the brain were not electrophysiologically connected, they were also found to be structurally not connected. Specifically, no streamlines were found between electrode sites that were electrophysiologically not connected (¬*E* → ¬*s*_1_), and these site pairs also did not intersect any known HCP white matter tract (¬*E* → ¬*s*_2_).

### 2.3 Structurally not connected electrode pairs lack “direct” electrophysiological connectivity

For *H*_3_ : ¬*S* → ¬*E*, only 40% of structurally not connected brain regions guarantee the lack of causal electrophysiological connectivity (see Figure 4B). We further analyze (and define) direct connectivity in the human brain. The heatmaps of time to *r*^∗^ for the neural features (F1, F2, F3) among not-connected pairs that are correctly labeled ¬*E* show that *r*^∗^ *<* 0.4 for F1 with time latency broadly dispersed, unlike the concentrated patterns of F1 in connected pairs. This suggests that the absence of a structural connection implies lower correlations with the F1 feature. Since the F1 neural feature is a result of an electrophysiological signal peak within 10 ms to 70 ms, it is compatible with a signature of direct electrophysiological signaling along structural pathways. Observing that the correlation with F1 is less than 0.4, we conclude that brain regions that are structurally not connected are not “directly” communicating electrophysiologically.

By contrast, the presence of F2 and F3 signal peaks after 70 ms in structurally not connected pairs suggests indirect or polysynaptic signaling. Removing electrode pairs suspected to be indirectly connected (*r*^∗^ *>* 0.4 for F2 or F3) from the analysis, we observe that the remaining structurally not connected electrode pairs are not directly connected electrophysiologically (*r*^∗^ ≤ 0.4 for F1) with 0.8 probability. Finally, since *H*_2_ is the contrapositive of *H*_3_, we observe similar results for *H*_2_ formalized as *P* (*E* → *S*) = *P* (*S* | *E*) which is 0.14 for both *s*_1_-connectivity and *s*_2_-connectivity. Relaxing the structural connectivity definition for *s*_1_-connectivity (*th*_s_1__ = 0) and for *s*_2_-connectivity (*th*_s_2 = 0 and non-zero overlap), we get *P* (*S* | *E*) equals to 0.3 and 0.6 respectively. We next tested *H*_2_ : *E* → *S* for “direct” connectivity by computing *P* (*S* | *E*) across increasing thresholds *th*_r_∗ on the *F* 1 feature (Table A3). As *th*_r_∗ increased, *P* (*S* | *E*) also increased for both structural definitions reaching ≈ 0.8. These results indicate that stronger correlations with the *F* 1 feature in the electrophysiological connectivity is increasingly associated with “direct” structural connectivity.

These results provide two important conclusions: (1) for future prospective studies, neurosurgeons can potentially place electrodes with ≈95% confidence between electrode sites that are known *a priori* to be structurally connected (*S* → *E*); (2) our tractography method leads to high true negative (¬*E* → ¬*S* and ¬*S* → ¬ “direct”*E*) confirming either indirect or no connectivity with ≈80% confidence when structural connection is absent.

### 2.4 Indirect structural connectivity implies higher latency in electrophysiological signaling

The F1 neural feature appears as a sharp wave (increased power in high gamma) with tight phase locking to the stimulation onset (strong ITPC) within the early stage after stimulation [16]. Previous study [16] showed that F1, defined in the time-frequency domain using power spectral density analysis, is comparable to the time-domain N1 component of the traditional CCEP signal and may serve as an indicator of feedforward connectivity. Our results above in Section 2.2 and 2.3 suggest that the presence of a structural connection is highly predictive of the presence of this early F1 feature in the evoked response. This supports the notion that F1 reflects direct electrophysiological signaling along structural pathways. By contrast, absence of structural connections imply lower correlation with F1 thereby suggesting more indirect or polysynaptic signaling.

We used a graph-theoretic shortest-path approach to identify indirect structural pathways involving an intermediate node. For understanding indirect structural connectivity and its relationship with delayed signaling, we constructed undirected electrode-to-electrode structural connectivity graph *G*^(s)^ = *V* ^(e)^*, E*^(s)^ and a directed graph *G*^(e)^ for causal electrophysiological connectivity for each of the ten subjects (see Methods for graph construction). In *G*^(s)^, an edge between two nodes exists when there are non-zero streamlines between them and an edge in *G*^(e)^ exists if the stimulation of one node evoked a significant response in the other electrode node. The graphs provide us a way to compute the sets of indirectly connected electrode pairs. We validate the indirect structurally connected electrode pairs from *G*^(s)^ in two ways: (a) must meet the directionality requirement from electrophysiological data (stimulating → recording) from *G*^(e)^, and (b) the expected time latency of the evoked signal in the indirect pathways must be atleast the sum of the latency in the direct pathways. For example, in the cortico-thalamo-cortical pathway, which we label as *C*1 → *T* → *C*2, we expect that the stimulation of *C*1 will evoke a response in *C*2 with a higher latency than the total latencies of indirect pathways through *T*. For the analysis that follows, we focus on four types of indirect pathways (visualized in order shown below in Figure 5A–D):

- Cortico-thalamo-cortical pathways (*C*1 → *T* → *C*2),
- Thalamo-cortico-cortical pathways (*T* → *C*1 → *C*2 or *C*1 → *C*2 → *T*),
- Thalamo-cortico-thalamic pathways (*T* 1 → *C* → *T* 2),
- Cortico-cortico-cortical pathways (*C*1 → *C*2 → *C*3).

**Fig. 5:**
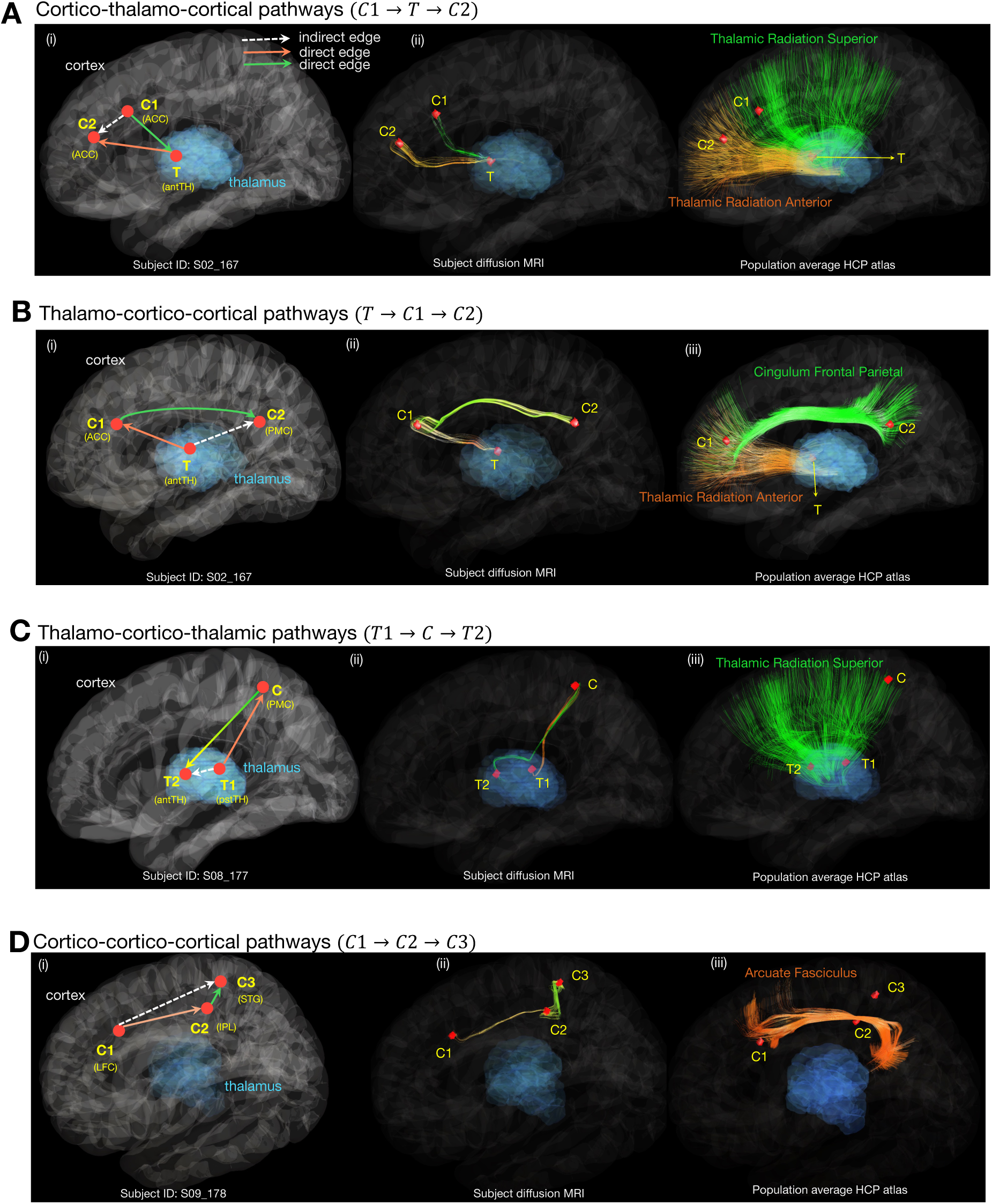
White matter pathways underlying indirect structural connectivity. The four types of indirect pathways are illustrated in **(A)** Cortico-thalamo-cortical pathways (*C*1 → *T* → *C*2) **(B)** Thalamo-cortico-cortical pathways (*T* → *C*1 → *C*2) **(C)** Thalamo-cortico-thalamic pathways (*T* → *C* → *T*) and **(D)** Cortico-cortico-cortical pathways (*C*1 → *C*2 → *C*3). (**A-D, i**) shows symbolic connections between three electrode sites. Solid lines (orange and green) indicate direct structural connections while dashed lines (white) indicate possible indirect structural connectivity when those electrode sites are not “directly” structurally connected from our *s*_1_ and *s*_2_ connectivity definition. The directionality of the causal electrophysiological connectivity (stimulation → recording) is used to validate the directionality of the indirect structural pathways. (**A-D, ii**) shows the corresponding streamlines from probabilistic tractography for the same three electrode sites (*s*_1_-connectivity). (**A-D, iii**) shows the white matter tracts from HCP tractography atlas that intersect the three electrode sites. The presence of indirect structural connections suggests possible polysynaptic pathways that may underlie the observed causal electrophysiological connectivity patterns. For example, in the cortico-thalamo-cortical pathway, we expect that the stimulation of *C*1 will evoke a response in *C*2 with a delay consistent with an indirect pathway through *T*. Time-to-first peak of the evoked signal is used to validate the time latency in the indirect pathways. STG: Superior Temporal Gyrus; IPL: Inferior Parietal Lobule; LFC: Lateral Frontal Cortex; ACC: Anterior Cingulate Cortex; PMC: Posteromedial Cortex; antTH: Anterior Thalamus; pstTH: Posterior Thalamus. We note that the *s*_2_-connectivity definition is more permissive, as it relies on electrode-tract intersection and will therefore miss indirect connections along the same tract and also when one of the sites lies in gray matter. Spatiotemporal videos depicting the propagation of causal electrophysiological signaling detected from individual electrode sites are provided in the Supplementary Information.

Note that direct pathways are electrode sites with a structural edge and an electrophysiological response between them in *G*^(s)^ and *G*^(e)^, respectively. Indirect pathways are endpoint pairs without a direct structural edge but connected through a shortest structural path with one intermediate node. By comparing the electrophysiological neural features (F1, F2, F3) of direct and indirect pathways from the graph objects, we observe that the most consistent effect was a reduction in the earliest component F1 for indirect relative to direct pathways (Figure 6). For cortico-thalamo-cortical signaling, the earliest component, *F* 1, was lower for indirect *C*1 → *T* → *C*2 pairs than for direct *C*1 → *C*2 pairs (mean = 0.18 versus 0.47). This was confirmed by a Mann–Whitney U test (*p* = 3.32 × 10^−9^) and was also evident in the raw CCEP signal shown in Figure 6A with no signal peak within 70 ms post-stimulation. Thalamo-cortical signaling showed the same pattern: *F* 1 was reduced for indirect *T* → *C*1 → *C*2 pairs compared to direct *T* → *C*2 pairs (mean = 0.33 versus 0.45, *p* = 2.28 × 10^−8^); see Figure 6B. For cortico-cortical signaling, *F* 1 was likewise lower for indirect *C*1 → *C*2 → *C*3 pairs than for direct *C*1 → *C*3 pairs (mean = 0.30 versus 0.47). This was confirmed by a Mann–Whitney U test (*p* = 8.96 × 10^−42^) and was also evident in the raw CCEP signal shown in Figure 6C. Thus, across all comparisons, indirect structural pathways were associated with weaker early electrophysiological responses. Mixed-effects model results for direct versus indirect pathway comparisons are reported in Appendix Table A4.

**Fig. 6:**
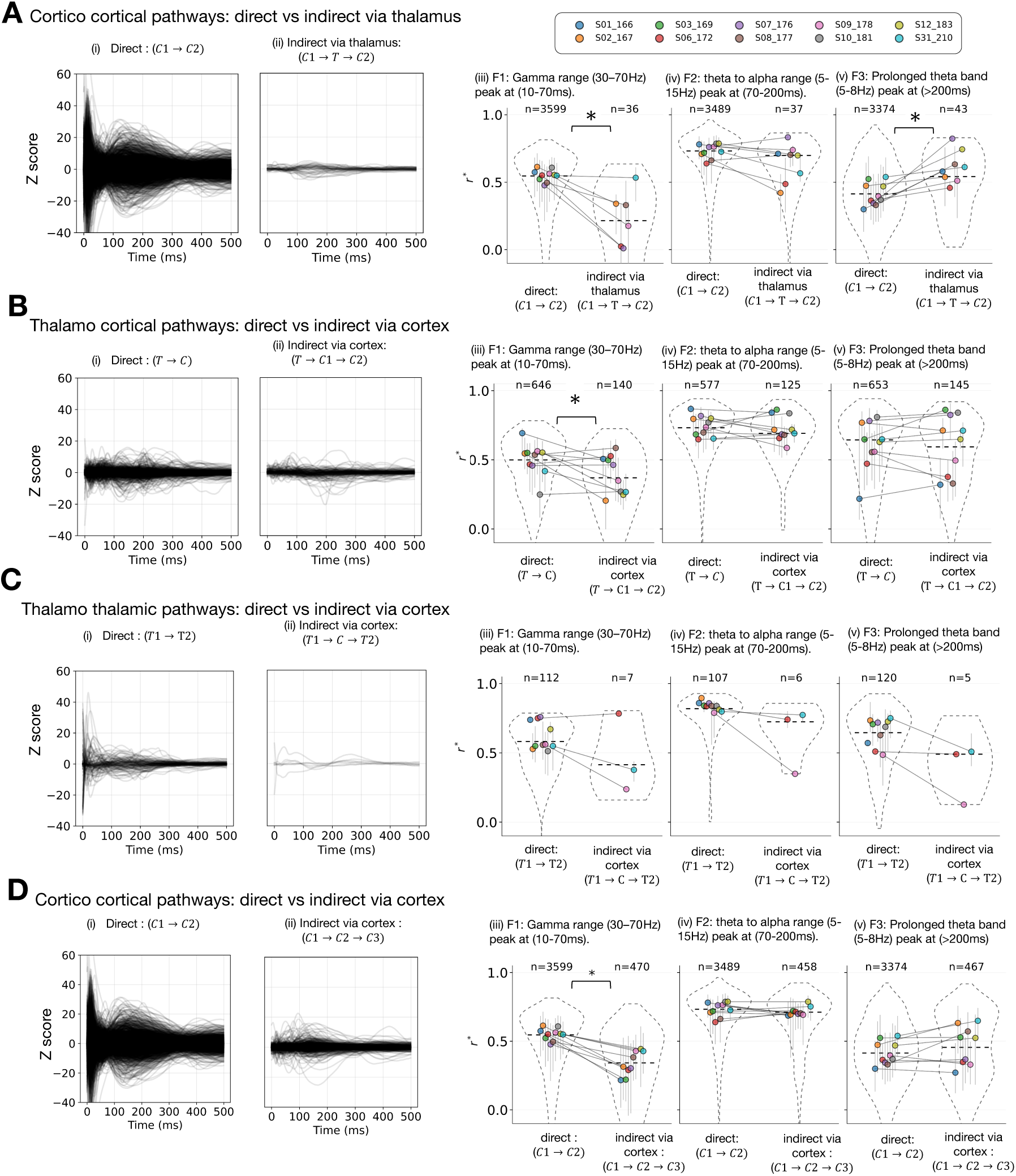
Matched CCEP and dMRI analysis distinguishes between direct and indirect connectivity and shows the involvement of thalamus in delayed oscillatory responses. **(A)** Corticocortical pathways with indirect connections via thalamus: (i) Baseline corrected and Z-scored raw signals for all direct-connections show peak response within 70 ms consistent with F1 feature while (ii) delayed response with peaks after 70 ms for indirect pathways. (iii) Correlation with F1 spectral features significantly reduce for indirect pathways relative to direct pathways. (iv) F2 spectral features do not differ significantly between direct and indirect pathways. (v) Increase in correlations with F3 spectral features for indirect pathways relative to direct pathways due to involvement of thalamus. **(B)** Thalamocortical pathways with indirect connections via cortex: (i) Baseline corrected and Z-scored raw signals for all direct connections show some peak response within 70 ms while (ii) delayed response with peaks after 70 ms for indirect pathways. (iii) Correlation with F1 spectral features significantly reduce for indirect pathways relative to direct pathways. (iv) F2 and F3 spectral features do not differ significantly between direct and indirect pathways. The presence of indirect structural connections suggests possible polysynaptic pathways that may underlie the observed causal electrophysiological connectivity patterns. **(C)** Thalamothalamic pathways with indirect connections via cortex: (i) Raw signals for all direct connections show peak response within 70 ms (ii) raw signals for indirect pathways with inconclusive number of data points. (iii, iv, v) No significant differences between direct and indirect connections. **(D)** Corticocortical pathways with indirect connections via cortex: (i) Raw signals for all direct connections show peak response within 70 ms consistent with F1 feature while (ii) delayed response with peaks after 70 ms for indirect pathways via cortex. (iii) Correlations with F1 spectral features significantly reduce for indirect pathways relative to direct pathways. (iv, v) F2 and F3 spectral features do not differ significantly between direct and indirect pathways. Significance in direct vs. indirect connections is indicated by asterisks (* *p <* 0.05, absolute difference in mean *>* 0.1). Each colored circle is the subject-wise median of peak correlations and vertical bars are the inter-quartile range.

The neural features that appear later, F2 and F3, changed less consistently. For *F* 2, the mean differences between direct and indirect pathways were small (*<* 0.1) in all three principal comparisons. For *F* 3, most direct versus indirect comparisons were again small, with one exception. The *C*1 → *T* → *C*2 indirect path showed a larger late component than direct *C*1 → *C*2 pairs (mean = 0.5 versus 0.4, with a p-value *p <* 0.05). These results indicate that the signature of indirect structural connectivity is attenuation of the earliest response component. Further, the F3 component is associated with connections involving the thalamus region. Together, these observations are consistent with the idea that indirect structural connectivity reflects multi-step propagation and therefore delayed electrophysiological signaling. For the small number of *T* 1 → *C* → *T* 2 pathways, none of the direct-indirect comparisons showed a robust difference in F1, F2, and F3 (see Figure 6D). This may also explain the structural pathway involved in the thalamo-thalamic connections and requires further study with denser thalamic stimulation and recording coverage. More examples of comparison between direct and indirect thalamic pathways are shown in Supplementary Figure A2 for *N* = 40 subjects under *s*_2_-connectivity definition.

For the *N* = 40 subjects analyzed under the *s*_2_-connectivity definition, we observed the same pattern, reduced F1 in indirect pathways relative to direct pathways, which further affirm the results presented above. We note that the *s*_2_-connectivity definition is less accurate than *s*_1_-connectivity at disambiguating between direct and indirect connections. This is possibly a result of tract intersection being more difficult to establish for electrode sites located in gray matter compared to tracking from the gray matter ROI (as in *s*_1_-definition); for example, in Figure 6D(iii), *C*3 is located in gray matter.

### 2.5 Diffusion microstructure properties correlate with conduction velocity

Conduction velocity is a critical neurophysiological property of brain function. It is well established that conduction velocity increases with axon diameter and myelination [32]. Multiple dMRI biophysical modeling methods [33–36] have been proposed to gain sensitivity to axon diameters and myelination, which could provide a proxy for conduction velocity. However, subject-specific validation of conduction velocity measurements is challenging. Here, we mapped dMRI-derived microstructure properties along the tracts of interest and determined their correlation with electrophysiological measurements of conduction velocity in the same subjects. This analysis helps to further elucidate the structure-function relationships that govern brain connectivity.

The conduction delay between the stimulated and recorded brain areas was estimated from the measured time-to-first peak of the CCEP (see Methods and illustration in Figure 7A, B). For *N* = 10 patients, tract length between electrode sites was estimated as the minimum of the streamline lengths between electrodes. We then defined the conduction velocity as the ratio of tract length and the time-to-first evoked peak from CCEP between a given pair of electrodes (see Figure 7A, and time-series in 7B). Microstructural features for each tract were obtained from (1) diffusion tensor fitting: fractional anisotropy (FA), mean diffusivity (MD) and (2) NODDI fitting [37]: neurite density (NDI), orientation dispersion (ODI), and free water fraction (FWF) as tract-weighted median (see Figure 7C). Within each patient, we computed correlations between conduction velocity and each microstructure metric using Pearson correlation for structurally connected electrode pairs with streamlines count greater than 100, that is, th_s_1 *>* 100 and present in the white matter, that is, non zero overlap with the white matter tracts defined by the HCP tractography atlas. Conduction velocity showed a positive correlation with FA and NDI and a negative correlation with MD and ODI while FWF showed non-significant correlations an individual patient level (see Figure 7D and Supplementary Table A5). For an example patient S02 167, the strongest association was observed for ODI, which showed a robust negative correlation with conduction velocity (*r* ≈ −0.53). FA showed a clear positive correlation (*r* ≈ 0.45), and both MD (*r* ≈ −0.37) and NDI (*r* ≈ 0.37) showed moderate correlations in the expected directions as shown in Figure 7D. Free water fraction (FWF) showed only a weaker positive association with conduction velocity (*r* ≈ 0.23). These results show that diffusion MRI microstructure properties can act as physiologically meaningful markers in quantifying the variation in speed of electrophysiological transmission in the brain. In order to demonstrate the importance of subject-specific dMRI and conduction velocity correlations, in an example patient S02 167 shown in Fig. 7E, diffusion microstructure features computed from the patient’s own dMRI showed stronger relationships with conduction velocity but when the same electrophysiological data were paired with dMRI from a different subject S01 166 acquired under the same protocol, these relationships weakened. While the trend in the correlations (positive for FA and negative for MD) remained same, Pearson correlation *r* for FA decreased from 0.45 to 0.3 and *r* for ODI decreased from -0.53 to -0.27.

**Fig. 7:**
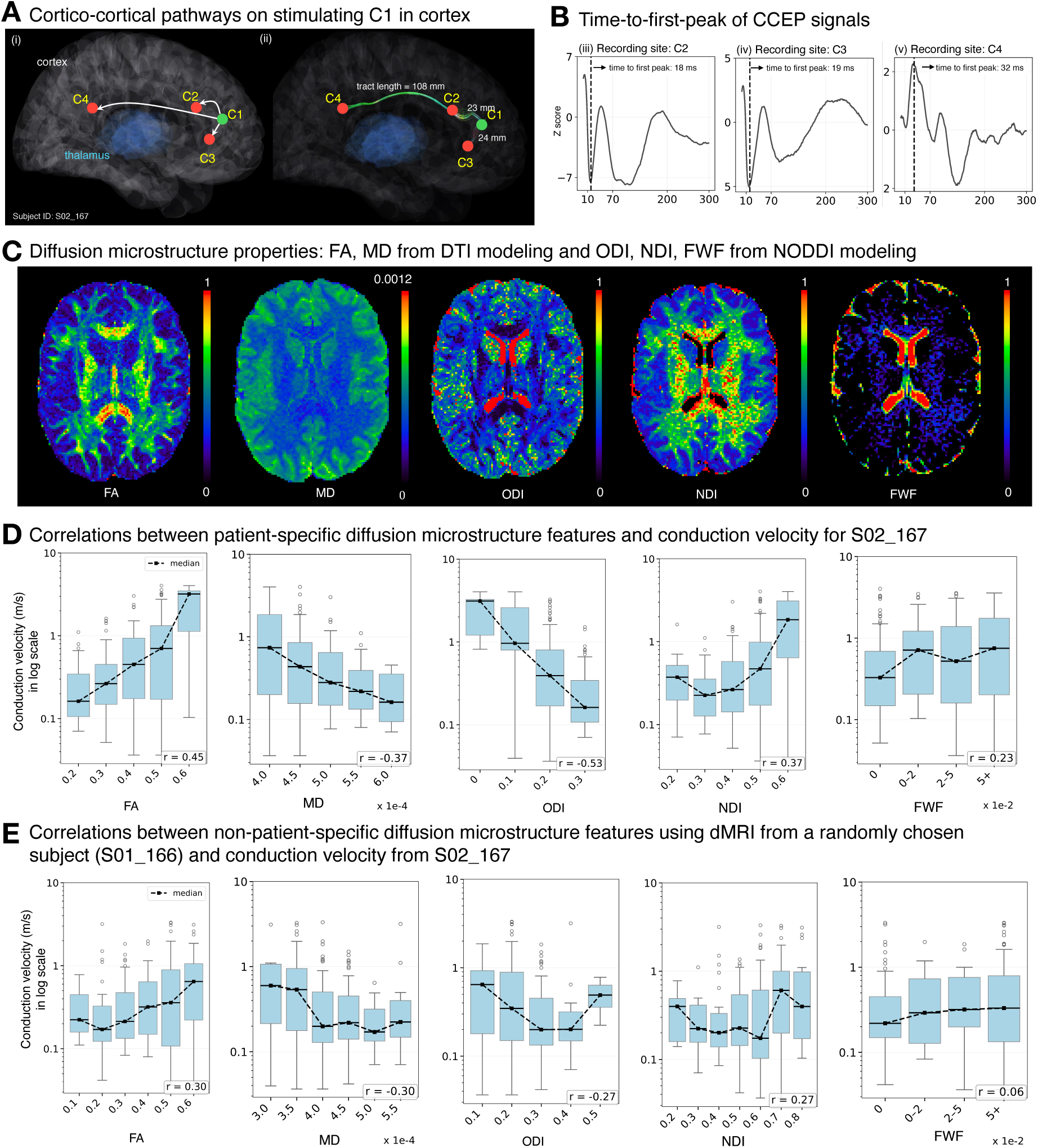
Conduction velocity and its correlations with patient-specific microstructure properties. **(A)** Cortico-cortical pathways on stimulating C1 electrode ROI in cortex: (i) electrode locations and recording directions mapped to MNI standard space and visualized using LeadDBS. (ii) reconstructed white matter tracts after probabilistic tracking for subject S02 167 with corresponding tract lengths. **(B)** Computation of time-to-first-peak on stimulating C1 electrode ROI in cortex and recording CCEP signals from (iii) C2, (iv) C3, and (v) C4. Vertical line denotes time-to-first-peak in each that is used in conduction velocity computation. **(C)** Computation of diffusion microstructure features: fractional anisotropy (FA) and mean diffusivity (MD) from dti fit using FSL, Orientation Dispersion Index (ODI), Neurite Density Index (NDI), and Free Water Fraction (FWF) from NODDI model fitting. **(D)** Correlations between patient-specific diffusion microstructure properties (FA, MD, ODI, NDI, FWF) and conduction velocity. We observe that conduction velocity positively correlates with FA and NDI and negatively with MD and ODI. These metrics are computed from patient-specific dMRI and correlated with conduction velocity computed using same patient’s CCEP signals. **(E)** Correlation between conduction velocity and diffusion microstructure properties using electrophysiological recordings from a (randomly chosen) patient: S02 167 and diffusion MR imaging from a different (randomly chosen) patient S01 166. Correlations with microstructure properties (FA, MD, ODI, NDI, FWF) are shown. Observe that the correlations are lower in this non-patient-specific case and the trends are much less pronounced. This demonstrates the importance of individual’s matched diffusion MRI in studying conduction velocity.

### 2.6 Pathways involving thalamus engage in dealyed oscillatory electrophysiological responses

Recently it has been shown that the stimulation of thalamic sites (i.e., anterior, mid, and posterior association nuclei) evokes electrophysiological responses in other thalamic sites [38]. To determine whether these evoked responses are mediated through cortical structures, we compared the spectral features of thalamo-thalamic (*T* 1 → *T* 2) and thalamo-cortico-thalamic (*T* 1 → *C* → *T* 2) pathways and found that there were no significant differences (Figure 6C for *s*_1_; Figure A2-C for *s*_2_). This is consistent with the idea that the thalamus participates in relaying electrophysiological signals across brain. Interestingly, we observe that electrodes in thalamo-cortical tracts showed increased conduction velocity compared to cortico-cortical tracts (Figure 8C). This is expected as thalamo-cortical tracts are known to be highly myelinated which explains the increase in conduction velocity. However, as discussed in Section 2.4, we found that involvement of thalamus in indirect structural pathways resulted in delayed latency (Figure 6A). We also compared the two indirect cortico-cortical pathways, one via cortex and the other via thalamus. While *F* 1 was generally low (below 0.5) for all indirect pathways we studied, cortico-thalamo-cortical (*C*1 → *T* → *C*2) pathways showed even lower *F* 1 correlations than cortico-cortico-cortical (*C*1 → *C*2 → *C*3) pathways. By contrast, *F* 3 correlations were higher for *C*1 → *T* → *C*2 than for *C*1 → *C*2 → *C*3, and this pattern was observed in 8 of 10 patients as shown in (Figure 8D). Also, using a linear mixed-effects model with pathway as a fixed effect and subject as a random intercept we observed F3 was significantly higher for *C*1 → *T* → *C*2 than for *C*1 → *C*2 → *C*3 (beta = 0.091, 95% CI [0.014, 0.167], p = 0.021; 484 observations from 8 subjects). These findings suggest that the thalamic involvement is associated with delayed electrophysiological signaling, with attenuation of earlier components and relative enhancement of later oscillatory F3 components. Together, this pattern suggests that signals involving the thalamus are not simply transmitted in a feed-through manner, but may additionally engage intra-thalamic processing or interactions before re-emerging along downstream pathways to engage the cortex in oscillatory theta band responses.

**Fig. 8:**
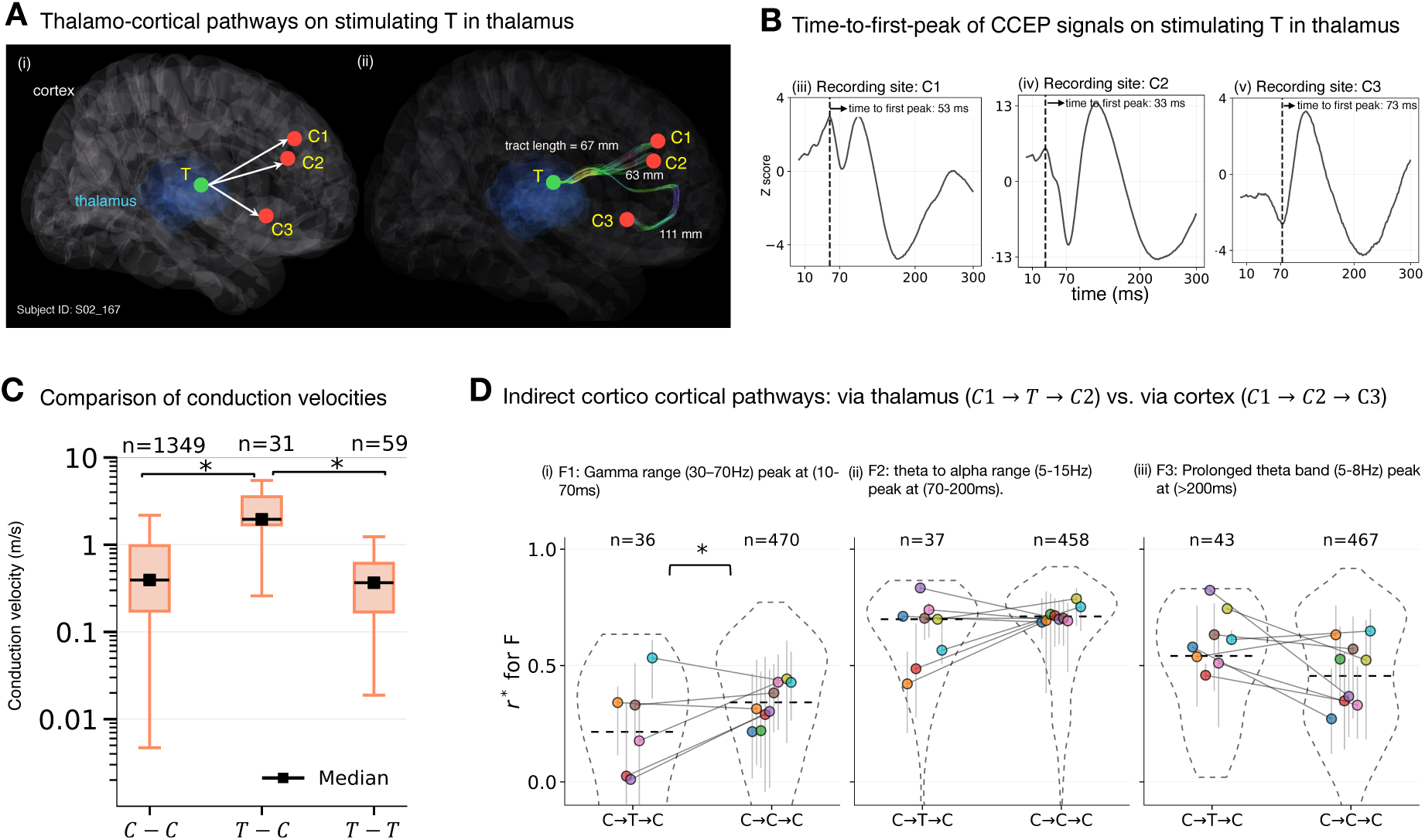
Thalamic involvement shifts indirect cortico-cortical signaling towards later electrophysiological components. **(A)** Thalamo-cortical pathways on stimulating T electrode ROI in thalamus: (i) electrode locations and recording directions mapped to MNI standard space and visualized using LeadDBS. (ii) reconstructed white matter tracts after probabilistic tracking for subject S02 167 with corresponding tract lengths. **(B)** Computation of time-to-first-peak on stimulating T electrode ROI in thalamus and recording from (iii) C1, (iv) C2, and (v) C3. Conduction velocity calculated as the ratio of white matter tract length to evoked response latency. Vertical line denotes time-to-first-peak in each that is used in conduction velocity computation. **(C)** Conduction velocity among thalamo-cortical (T-C) pathways is substantially higher than the cortico-cortical (C-C) or thalamo-thalamic (T-T) pathways. **(D)** Comparison of two indirect pathways for (*N* = 10): via the thalamus (*C*1 → *T* → *C*2) and via an intermediate cortical node (*C*1 → *C*2 → *C*3). Subject-level and pooled comparisons of the three spectral CCEP features: **(i)** *F* 1, gamma band (30–70 Hz), peaking at 10–70 ms; **(ii)** *F* 2, theta-to-alpha band (5–15 Hz), peaking at 70–200 ms; and **(iii)** *F* 3, prolonged theta band (5–8 Hz), peaking at *>* 200 ms. Dashed violins show pooled distributions, horizontal dashed lines indicate pooled medians, and colored points connected across conditions show subject-wise medians with interquartile ranges. Sample sizes (*n*) denote the number of electrode pairs in each pathway class. *C*1 → *T* → *C*2 pathways showed lower *F* 1 and higher *F* 3 than *C*1 → *C*2 → *C*3, consistent with attenuation of earlier components and relative enhancement of later components when the thalamus is involved in indirect signaling.

## 3 Discussion

While it is well-understood that white matter fiber bundles are the structural conduits of information flow in the brain, here we address how this structural network constrains causal electrophysiological signaling. We predict causal electrophysiological connectivity between pairs of brain sites using diffusion MRI tractography. We show how streamline counts or probability thresholds of intersection with white matter and spectral analysis of electrophysiological signals can provide tunable confidence levels to study the relationships between structural and electrophysiological connectivity. Our findings are consistent with recent efforts towards linking structural and functional connectivity in the human brain [24, 39–41]. However, our study leverages unique multi-modal data to achieve several distinct contributions that differentiate it from prior work and establish new capabilities for designing and targeting neuromodulatory treatments.

### White matter pathways explain the spectral signature of electrophysiological signaling

To provide insights into the structural pathways underlying causal electrophysiological signaling, we complemented the SEEG recording with dMRI tractography. Across more than 150,000 stimulation-recording pairs, the presence of a tractography-defined pathway predicted electrophysiological connectivity with high probability (∼ 0.95), a result replicated using both subject-specific tractography and an independent population atlas. These findings extend prior studies linking anatomy to spontaneous functional correlations by showing a substantially tighter correspondence when connectivity is assessed causally rather than inferred from ongoing activity [18]. In this context, our approach showed that white matter architecture is not simply correlated with functional coupling but predicts the capacity for directed signal transmission between brain regions. Thus, our approach provides a higher spatial resolution mapping to guide the placement of electrodes in functional neurosurgeries such as SEEG for epilepsy treatment and deep brain stimulation for neuropsychiatric disorders.

Our multi-modal SEEG and dMRI tractography data rendered us uniquely positioned to disambiguate between direct and indirect connectivity. Recent work showed that evoked responses can be grouped into three spectral components: F1, F2, and F3 reflecting early or late phase coherence and peaks across different frequency bands [16]. We characterized the structural substrates underlying distinct signatures of stimulation-evoked responses to direct versus indirect cortical and thalamic white matter pathways. In fact, the most robust signature of direct communication was the early F1 component, a phase-locked broadband response within 10-70 ms that was consistently enriched among structurally connected pairs and attenuated in indirect pathways. By contrast, the most delayed component (F3) was a more prominent oscillatory theta rhythm whenever thalamic sites were involved in direct or indirect pathways. These observations provide unique temporal information about the hierarchy and sequence of communication in which early responses index rapid transmission along relatively direct axonal routes (including cortico-thalamic direct routes), whereas later responses may reflect recurrent network engagement, information integration with a significant delay at the level of thalamus.

### Involvement of thalamus in delayed and oscillatory electrophysiological responses

Our work leverages the unprecedented intra-thalamic recordings in the human brain with thalamic data providing particularly informative and novel lines of information. It is well known that early components in CCEP signals are correlated with direct structural connections [24, 25]. This also leads to a natural presumption that later spectral components in CCEP signals would imply indirect connectivity. However, we find that the role of thalamus is key in studying indirect connectivity and distinguishing the spectral signatures in CCEP signals. Our results show that even for directly connected brain regions, if the thalamus is involved, both early components (F1 feature) and delayed components in the theta range (F3 feature) are present. This counters the assumption that direct connectivity is only correlated with rapid high-frequency activity immediately after stimulation. Further, on analyzing all possible topologies of indirect connectivity, we find a low association with the F1 feature and a presence of the F2 feature. But when thalamus is involved in these indirect connections, the F3 feature is additionally prominent.

Recent theoretical models discuss the role of thalamus as a transformer or a gate for signals reentering into the cortex [42–45]. To provide quantitative evidence for this, we identified all the directly connected electrode sites and those that were indirectly connected via cortical (e.g., *C* → *C* → *C*) or thalamic nodes (*C* → *T* → *C*). Interestingly, we show that cortico-thalamic tracts exhibited significantly faster conduction velocities than many cortico-cortical pathways. But, simultaneously, the involvement of thalamus within indirect routes was associated with significant delayed downstream responses and enhanced late oscillatory activity (F3). This provides direct evidence in support of thalamus acting as a transformer for signals reentering into the cortex. Further, the combination of rapid thalamo-cortical transmission with delayed network-level consequences may reflect distinct feedforward and recurrent functions operating on different timescales.

### Patient-specific diffusion MRI is important for estimating conduction velocity

Beyond the binary definitions of connectivity, we also investigated the temporal components of connectivity in CCEP signals and how these correlate with microstructure properties computed using diffusion MRI. It is well established that conduction velocity increases with myelination [32]. Therefore, there has been a longstanding interest to map myelination along white matter tracts using non-invasive MRI methods and use this to estimate conduction velocity along specific tracts. Previously, this topic has been studied using large-cohort CCEP resources such as F-TRACT [26] and HCP database acquired in separate set of subjects. Our study is unique because we acquire high-resolution dMRI and CCEP data in the same patients who underwent SEEG at our hospital. This gives us a rich dataset to study direct structure-function correlations in the same brains. Our results show that, for a given subject, diffusion microstructure properties measured along personalized tractography-defined pathways, such as higher FA and NDI and lower MD and ODI, correspond to faster conduction velocity. While these microstructure properties represent parameters of simplistic models, we know that FA increases with increasing myelin content and MD follows the reverse trend [46]. Still, these measures lack specificity and are also affected by other factors. However, these results do support the expected correlation that pathways with more myelination conduct more rapidly and have increased FA, whereas pathways with less myelination have lower FA and greater MD and conduct more slowly. This interpretation is consistent with prior observations from deep brain stimulation studies, in which evoked potential amplitude correlated with FA and tract volume, while response delay correlated with tract length, together suggesting that the magnitude and timing of DBS-evoked effects are shaped by the integrity and geometry of the underlying white matter pathways [47].

We note an important distinction between using patient-normative diffusion MRI data to identify whether an electrophysiological connection is structurally supported versus estimating/correlating conduction velocity. We observed that *s*_1_-connectivity results were similar to *s*_2_-connectivity results and also similar to using averaged streamline information from 200 HCP subjects (see methods in Section 4.8 and Figure A1). This suggests that patient-normative diffusion data can be utilized to predict electrophysiological connectivity. Interestingly, this did not hold for conduction velocity. In the example patient shown in Figure 7D, E, diffusion microstructure features computed from the patient’s own dMRI showed stronger relationships with conduction velocity. When the same electrophysiological data were paired with dMRI from a different subject acquired under the same protocol, these relationships weakened. Thus, while structural connectivity estimates appear relatively robust to the use of non-specific diffusion data, correlation of conduction velocity with microstructure properties remain patient-specific for reliable interpretation.

### Limitations and technical considerations

Due to the limitations of tractography and the SEEG data acquired from the epilepsy patients, the results presented here also have intertwined limitations of both modalities. Firstly, accuracy of tractography methods depends on many aspects such as acquisition, preprocessing, co-registration, fiber-orientation modeling, and seed definition. As such, there is a lack of standardization which may contribute to different levels of anatomical precision. The geometry of a white matter tract can make it easier or more difficult to map with tractography relative to other tracts [20, 21]. Further, many different models of intra-voxel crossing fibers exist and the choice of model also influences tractography results. Still, it is well established that the dominant orientation(s) in white matter corresponds to the local orientation of white matter tracts. Here, we utilized a well-established, multi-directional fiber model [48, 49] to perform tractography rigorously to a standard that correlates with electrophysiological function. Indeed, our findings provide a new level of functional interpretation of the structural tractography methods across the cortical and subcortical structures at a tunable confidence. Secondly, the electrophysiological signaling from epilepsy patients may not be representative of healthy individuals. However, we note that 80% of the recordings in the data that we used were done in non-epileptic and healthy tissue [50–52] in the brain. So only a minority of SEEG recordings are from epileptic tissue. Further, across subjects, since the epileptic tissue is not the same, the epileptic (minority) recording sites together with healthy (majority) sites across patients mitigate the possibility of a systematic bias affecting our measurements. Although chronic epilepsy involves molecular, cellular, and local circuit changes, currently there is no firm evidence that indicates a whole-brain reorganization of large-scale network topology in these patients relative to healthy controls. Finally, we note that it is impossible to get this data from healthy individuals given the invasive nature of the procedure.

## Conclusion

In closing, we foresee our findings to have both scientific and translational impacts. Scientifically, they establish a multiscale framework linking anatomical pathways, microstructure, and causal dynamics in the living human brain. Translationally, our models can improve targeting for invasive monitoring, optimize stimulation therapies, and help predict how lesions or white matter degeneration alter network communication. Progress toward clinical translation will require standardized imaging protocols and automated analysis pipelines of the kind we present here. Especially with the use of SEEG extending to neuropsychiatric patients — we envision a non-invasive tool for guiding electrode placement and helping with the interpretation of results. Since the circuitry involved in many neuropsychiatric conditions is not yet well understood and many new neuromodulation treatments such as low-intensity focused ultrasound are rapidly developing, our framework provides a systematic way to understand the underlying circuitry and predict signaling behaviors. More broadly, our results suggest that the human connectome is best understood as a temporally organized communication system in which anatomy constrains the flow of information.

## 4 Methods

### 4.1 Participants

We conducted a retrospective study of 40 patients with medically refractory epilepsy (40% female; mean age ± SD: 37.9 ± 12.3) who underwent a SEEG procedure (Table A1). All participants underwent minimally-invasive electrophysiological monitoring at Stanford Hospital as part of their standard clinical care for refractory epilepsy. On average, 143 ± 33 (mean ± SD) electrode sites were implanted per participant. The entire dataset encompasses 5794 brain sites across all subjects, with each adjacent pair stimulated approximately ∼45 times (with sufficient inter-trial intervals to avoid overlapping effects), while recordings were taken from every other electrode sites. Post-implant CT scan and pulse-evoked potentials (*N* = 40) were acquired during the SEEG for clinical care without any input from us.

### 4.2 Patient safety and ethics

All procedures involving human subjects were conducted with approval from the IRB of Stanford University. Informed consent was obtained from all participants prior to enrollment in the study. The electrode implantation strategy was determined by a multidisciplinary team of clinicians based on the patient’s clinical needs and was not influenced by this research project.

### 4.3 DWI acquisition, preprocessing, and microstructure properties

Out of 40 patients in our study, 10 of them volunteered for diffusion MRI scan. From Table A1, ten patients for whom we have matched diffusion scans have asterisk (∗). Multi-shell diffusion data with 1.5 mm isotropic resolution were acquired prior to the SEEG procedure of each patient using a 2D single-shot diffusion-weighted spin echo (DW-SE) echo-planar imaging (EPI) on a 3T GE Premier MRI. Five “blip up” b=0 and 1 additional “blip down” b=0s were collected for distortion and motion correction. Diffusion pulse sequence parameters are listed in Figure 2C and the processing pipeline is outlined in Figure 2E. Diffusion-weighted imaging data were first converted from dicom to `nifti` (Neuroimaging Informatics Technology Initiative) format using dcm2niix. Susceptibility and eddy current-induced distortion estimations and corrections were performed using FSL’s topup and eddy cuda10.2. Diffusion tensor fitting was then performed using `dtifit` to generate maps of fractional anisotropy, mean diffusivity, and principal eigenvectors for quality checks. To model fiber orientations in diffusion MRI data, we applied bedpostx gpu. This method estimates the posterior distribution of diffusion parameters using Markov Chain Monte Carlo (MCMC) sampling. Diffusion Tensor Imaging (DTI) fitting was performed using `dtifit` to generate maps of fractional anisotropy (FA) and mean diffusivity (MD). FA provides a measure of the directional coherence of water diffusion, while MD reflects the overall magnitude of diffusion.

### 4.4 Co-localization of electrodes

Post-surgery CT scans were acquired after electrode implantation. Precise electrode positions were determined in FreeSurfer surface-space, voxel-space, and MNI-space by the iElVis toolbox [53]. The post-implant CT scan was co-registered to the pre-implant T1-weighted MRI using the `flirt` function from the Oxford Centre for Functional MRI of the Brain Software Library or using `bbregister` from Freesurfer [54] to get the best results. The electrode coordinates in the native anatomical space were inspected and manually labeled on the T1-registered CT image using BioImageSuite [55] by an experienced neurologist (JP), based on the individual morphology of the brain and the associated landmarks. A 2 mm sphere centered on the centroid of each bipolar electrode site was used to define the anatomical location and for seeds and targets in tractography. We co-registered the electrode sites by registering the native T1-weighted image to the *b* = 0 image for analysis in the subject’s diffusion space and co-registered to MNI space for analysis in standard MNI space (see Figure 3B) using the ANTs library package.

### 4.5 Causal electrophysiological connectivity

SEEG involves implanting ∼150 electrodes deep in the brain and is used routinely for medically refractory epilepsy patients to determine the seizure onset zone and seizure propagation patterns. During the SEEG procedure, repeated single electrophysiological pulse stimulation is used to study causal electrophysiological relationships in the brain. Each CCEP trial consisted of a single stimulation of an instantaneous (pulse duration = 0.2 ms) 6 mA biphasic square wave pulse repeated (∼ 45 trials) in bi-polar manner. To determine causal electrophysiological connectivity during SEEG, we used a previously reported semi-supervised learning algorithm based on the spectrogram of evoked potentials across time and frequencies [16]. In this approach, the data was first partially and tentatively labeled based on two measures of connectivity: (1) trial-averaged power of evoked potentials as a measure of strength, and (2) inter-trial phase coherence given by

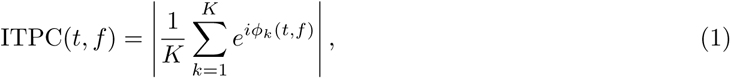

across repeated stimulation as a measure of consistency for the evoked responses.

A semi-supervised machine learning approach (UMAP [56]) was applied to learn the inherent data structure from this preliminary labeling to re-classify the data within each subject into two separate clusters: one cluster of data where stimulation of the seed site caused significant evoked potentials in the target site, and another cluster of data where stimulation of the seed site did not cause any significant evoked potentials in the target site. These two clusters formed the electrophysiological “connected” and “not-connected” labels, respectively in our study.

This also resulted in the significant time/frequency boundaries of three distinct neural features in the electrophysiology data:

- F1: high-frequency (30 − 70 Hz) peak power immediately after the stimulation (10 − 70 ms).
- F2: theta to alpha range (5 − 15 Hz) peak power at ∼ 120 ms post-stimulation (70 − 200 ms).
- F3: prolonged (*>* 200 ms) power increase in the theta band (5 − 8 Hz) with oscillation.

The output connectivity predictions of this machine learning method are then verified and augmented with human expert’s (neurosurgeon) input. This approach is based on threshold-agnostic measurements of changes in the power spectrum and it relies on the measure of inter-trial phase coherence (ITPC [42]) to ensure that only responses that are consistently evoked across trials are included. Using a sliding-window correlation approach, the maximum strength of these feature representations (Pearson correlation: *r*∗) and their presence in time are utilized to distinguish direct versus indirect electrophysiological connections. In this paper, we study how distinct white matter pathways correspond to these three neural features: F1, F2, and F3, as described above.

### 4.6 Structural connectivity: *s*_1_ and *s*_2_

We constructed a 2 mm sphere as a region of interest (ROI) around the centroid of each bipolar electrode sites and proposed two definitions of structural connectivity (see Figure 3):

- *s*_1_-connected (for *N* = 10 patients): We performed probabilistic tractography for individual subject. A pair of stimulating and recording electrodes is *s*_1_-connected if the total number of streamlines between them is greater than a threshold called th_s_1. Probabilistic tractography in the native diffusion space of the subject was performed using `probtrackx2`. Seeds were placed in the 2 mm ROIs and tracking was constrained using target masks, waypoint masks, and stopping criteria.
- *s*_2_-connected (for *N* = 40 patients): We use a population-based probabilistic tractography atlas that aggregates voxel-wise white matter tract probabilities from 1,065 HCP young adult tractograms generated via automatic and augmented fiber tracking. Each native T1-weighted image of the subject was registered to the ICBM 2009a nonlinear asymmetric T1 template, and the resulting transform was applied to the electrode ROIs to co-register them in MNI space. We constructed atlas-defined tract mask with probability exceeding th_s_2 and computed its overlap with the ROI of electrodes. Here, we only study electrode ROIs that have non-zero overlap with the white matter. We call a pair of electrodes *s*_2_-connected if both are located in the same white matter tract from the HCP tractography atlas [29] above a certain probability (th_s_2) and at least 50% of ROI overlaps with the tract.

Causal electrophysiological connectivity is unidirectional while structural connectivity is not. Therefore, we labeled pairs of bipolar sites as electrophysiologically connected if either direction is electrophysiologically connected. Pairs not connected in both directions were labeled as electrophysiologically not connected. When a pair of electrodes were electrophysiologically not connected and the testing of the causal electrophysiological connectivity was only done in one direction, we excluded such pairs from the analysis since the other direction was not tested. Total number of electrode pairs labeled as electrophysiologically connected (*E*) and not connected (¬*E*) can be found in the Table A1.

### 4.7 Tunable parameters of the connectivity framework

Our framework contains tunable parameters at both the electrophysiological and structural connectivity, allowing the connectivity definition to be adapted to the intended use case. Electrophysiological connectivity is controlled by the feature choice (*F* 1, *F* 2, or *F* 3) and by the correlation threshold *th*_r_∗. Increasing *th*_r_∗ yields a more stringent definition of connectivity, whereas lower values provide a more relaxed definition. We hypothesize that *F* 1 depicts direct structural connectivity and is therefore the preferred feature when the goal is to identify likely direct pathways. We use *r*^∗^ *>* 0.4 to show the presence or absence of F1-3 features.

Structural connectivity is based on the *s*_1_- and *s*_2_-connectivity definitions. For *s*_1_ definition, *th*_s_1__ = 0 corresponds to the most permissive setting, in which any non-zero streamline count after probabilistic tractography is considered evidence of connectivity. For *s*_2_ definition, connectivity depends on both the tract threshold *th*_s_2 and the overlap of electrode ROI with the tract region from the HCP atlas. Under the *s*_2_-connectivity definition, a pair of stimulating and recording electrodes was classified as structurally connected if both electrode ROIs overlapped the same white matter tract from the HCP tractography atlas above a tract probability threshold *th*_s_2 and a minimum overlap-area threshold of 50%. In this formulation, *th*_s_2 tunes the confidence of tract assignment, while the overlap threshold tunes the required extent of electrode–tract intersection. Smaller thresholds provide a more relaxed definition of connectivity, whereas larger thresholds provide a more conservative definition. Thus, the framework can be tuned according to the intended use case, ranging from broad network-level inference to higher-confidence identification of direct pathways.

### 4.8 HCP-based structural connectivity

An alternate definition of structural connectivity which lies in between *s*_1_-connectivity and *s*_2_-connectivity is the structural connectivity definition using the HCP data. This definition has the advantages of probabilistic tractography but does not require MRI data acquired on each patient. In this approach, we used 200 randomly sampled HCP subjects to get a population-averaged estimate of streamline counts between electrode pairs. Diffusion imaging protocol in the HCP dataset is different from our patient dataset. This is because the HCP data has a higher spatial resolution of 1.25 mm isotropic and 270 diffusion directions across three shells of b=1000, 2000, and 3000 s mm^−2^. We used the same tractography parameters for both datasets to ensure consistency in streamline generation. List of HCP subjects in this study are listed in the Supplementary Table A2. In this study, 200 HCP participants (89 M, 111 F; mean age = 29 ± 3.5 years) were included. The T1-weighted scan of the example patient (S09 178) was registered to each HCP subject’s distortion-corrected diffusion space and the resulting affine-transform was applied to each 2 mm ROI around the electrode. Diffusion scan from each subject was preprocessed using the similar preprocessing steps as described in Figure 2E (top up, eddy current, and bedpostx followed by probtrackx2). The streamline count between a given electrode pair was quantified by averaging across the 200 HCP subjects to get a proxy streamline count for that electrode pair.

### 4.9 Indirect structural pathways

We defined two types of structural connection: direct or indirect. A pair of electrodes is defined as directly connected when probabilistic tractography identified streamlines between them above a specified threshold under the *s*_1_-connectivity definition. A pair of electrodes is defined as indirectly connected when a direct connection was absent but the connection was mediated through an intermediate node. Under the *s*_2_-connectivity definition, we similarly defined a structural pathway between a stimulation and recording electrode as direct when both sites were linked through a single white matter tract from HCP atlas, and indirect when the connection traversed more than one tract.

To formally define and compute direct and indirect connectivity, we constructed undirected electrode-to-electrode structural connectivity graph *G*^(s)^ = *V* ^(e)^*, E*^(s)^ for each subject (*N* = 10). Here, each bipolar electrode contact was represented as a node and an edge was added between two nodes when the streamline counts (after `probtrackx2`) between them exceeded a threshold (th_s_1 = 0). We also constructed a directed graph *G*^(e)^ for causal electrophysiological connectivity in the same subject where an edge from node *i* to node *j* was added if the stimulation of electrode *i* evoked a significant response in electrode *j* (i.e., the UMAP classification label for the pair is equal to 1, or activated [16]). Given that the precise anatomical boundaries of the thalamic nuclei within each participant may be variable and difficult to ascertain with neuroimaging data, and that the field of evoked responses or electrical stimulation may cross the nuclear boundaries, we refrained from labeling thalamic sites according to specific nuclei. Instead, we referred to them as anterior thalamus (antTH), middle thalamus (midTH) and posterior thalamus (pstTH). Contacts labeled as antTH, midTH, or pstTH were jointly classified as thalamic (*T*), whereas cortical (*C*) contacts were defined as sites outside the thalamus, amygdala, hippocampus, and basal ganglia. We then classified electrode pairs into direct and indirect pathways. Thalamo-thalamic edges were retained only when both contacts were thalamic and on the same side of the brain (that is, ipsilateral sites). These were difficult to track under probabilistic tractography used in *s*_1_-connectivity due to poor SNR.

Direct pathways were structural edges in *G*^(s)^ that also had a directed electrophysiological edge between the same two nodes *G*^(e)^. The direct pathway analyzed here were *C* → *C*, *T* → *C*, and *T* → *T*. Indirect pathways were shortest structural paths of length 2, that is, paths with one intermediate node in *G*^(s)^. We confirmed the indirect structural pathways by validating using two independent measures: (1) **directional consistency**: from the electrophysiological data we ensure that the directionality, stimulating → recording, from *G*^(e)^ is consistent in the observed pathway, (2) **latency consistency**: this is a criterion for latency in the indirect propagation, relative to direct propagation. That is, any indirect structural pathway must satisfy

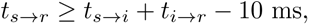

where *t*_s→r_ denotes the latency of the evoked response from the stimulation site *s* to the recording site *r*, *t*_s→i_ denotes the latency from the stimulation site *s* to the intermediate site *i*, and *t*_i→r_ denotes the latency from the intermediate site *i* to the recording site *r*. This criterion ensures that the endpoint latency of an indirect pathway is consistent with multi-step signal propagation, allowing for a 10 ms tolerance to account for measurement variability and uncertainty in peak detection.

The indirect pathways were classified into four categories: (1) cortico-cortico-cortical (*C*1 → *C*2 → *C*3), (2) cortico-thalamo-cortical (*C*1 → *T* → *C*2), (3) thalamo-cortico-cortical (*T* 1 → *C*1 → *C*2), and (4) thalamo-cortico-thalamic (*T* 1 → *C* → *T* 2) based on their labeled anatomical locations. We investigated how the spectral features of causal electrophysiological connectivity vary with respect to direct and indirect structural pathways. Peak correlation (*r*^*^) for each spectral feature F1, F2, F3 were statistically compared between direct and indirect pathways using a Mann–Whitney U test. The raw signals were simultaneously qualitatively compared after the artifact window (*<* 10 ms) and removing the labeled bad channels contaminated by stimulation artifact. For this comparison, we also excluded the recording bipolar sites that were too close to the stimulation site. That is, if the Euclidean distance between the centroid points of the two electrodes was *<* 5 mm then it was excluded from the analysis. For each pathway comparison and electrophysiological feature (*F* 1, *F* 2, *F* 3), we compared direct and indirect pathway distributions using two-sided Mann–Whitney U tests. Effect sizes were quantified using Cohen’s *d*. To account for repeated observations within subjects, we additionally fit linear mixed-effects models with a fixed effect for pathway type and a subject-level random intercept. Subjects were included in a given comparison only if they contributed at least one observation to both pathway groups. P-values were corrected for multiple comparisons using the Benjamini–Hochberg false discovery rate procedure.

### 4.10 Estimation of streamline count-weighted microstructure features

To further characterize the microstructural properties of white matter tracts between electrodes, we performed Neurite Orientation Dispersion and Density Imaging (NODDI) [37] fitting using the AMICO toolbox [57]. We use the default parameters for the NODDI model: parallel diffusivity *d*_||_ = 1.7 × 10^−3^ mm^2^*/*s and isotropic diffusivity *d*_iso_ = 3.0 × 10^−3^ mm^2^*/*s. This method estimates parameters such as the orientation dispersion index (ODI), neurite density index (NDI), and free water fraction (FWF) from multi-shell diffusion data. These parameters provide insights into the complexity and density of neurite structures within the brain, which relates to causal electrophysiological connectivity patterns observed in SEEG. We derived tract-level microstructural features, fractional aniosotropy (FA) and mean diffusivity (MD), from diffusion tensor fitting. For each electrode pair (*s, r*), FSL probtrackx2 was used to generate a voxel-wise probabilistic tractography map *W*_sr_, which served as a streamline count weight map over the voxel grid Ω. To summarize the microstructure along each tract, we computed streamline count-weighted statistics of each voxel-wise map *X*^(m)^, with *m* ∈ {FA, MD, FWF, NDI, ODI}. To emphasize the tract core, weights were thresholded at the 75th percentile of the nonzero values in (*W*_sr_). Weighted medians of the microstructural features were then computed across voxels with positive weights and finite map values. We correlated these microstructure metrics for structurally connected electrode pairs with the conduction velocity of the evoked potentials to understand how microstructural properties relate to the speed of neural signal transmission.

### 4.11 Conduction velocity

To estimate the conduction velocity of evoked potentials, the latency *t*_latency_ was defined as the time from the stimulation pulse to the first significant peak in the evoked potential that exceeded a predefined threshold based on baseline activity and after 10 ms artifact window (see Figure 6). We then calculated the geodesic distance as the tract length using the dipy Python package. Tract length of streamlines were calculated from streamline length distributions between electrode sites, with the minimum representing the shortest anatomically-valid pathway after filtering for connectivity between dilated electrode regions denoted by *L*_tract_. We then defined conduction velocity CV as:

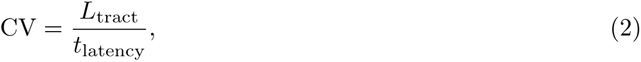

that is, the ratio of tract length and the time-to-first evoked peak from CCEP between a given pair of electrodes gives us CV. This yields a measure of how quickly neural signals propagate along the structural pathways connecting the stimulated and recorded sites. We correlated these conduction velocity estimates with microstructural properties of white matter tracts (FA, MD, ODI, NDI) to investigate how microstructural integrity relates to signal transmission speed in structurally connected electrode pairs. Conduction velocity was computed for electrode pairs with nonzero overlap with white matter.

## Data availability

Processed structural connectivity data from diffusion MRI and causal electrophysiological connectivity data for replicating the results will be made available upon publication. HCP probabilistic atlas data is publicly available at https://brain.labsolver.org/hcp_trk_atlas.html. HCP diffusion MRI data for 200 subjects is publicly available at https://db.humanconnectome.org/data/projects/HCP1200/. The list of HCP subjects information are provided with this paper. The thalamic atlas [58] used for visualization is available publicly on LeadDBS at https://www.lead-dbs.org/helpsupport/knowledge-base/atlasesresources/.

## Code availability

All code and data for design and implementation of the framework will be available upon publication.

## Acknowledgment

S.S. discloses support for the research of this work from Wu Tsai Neuroscience Institute Postdoctoral Fellowship at Stanford University, E.D. discloses support from NIH K99AG080076, and J.A.M. discloses support from NIH R01NS095985 and General Electric Healthcare. We thank Babak Razavi (Stanford University, USA), and Joanne Lau (Stanford University, USA) for their assistance with subject recruitment. We thank Sofia Pantis (Stanford University, USA) for her help with the data management and sharing. We sincerely thank the patients who participated in this study and the epilepsy care team. Saad Jbabdi (Oxford University, UK) and Paul McCarthy (Oxford University, UK) provided the code for streamlines visualization after probabilistic tractography. We thank Alexander Atalay (Stanford University, USA) for his help with the creation of videos for the electrophysiological signal propagation. Data were provided [in part] by the Human Connectome Project, WU-Minn Consortium (Principal Investigators: David Van Essen and Kamil Ugurbil; 1U54MH091657) funded by the 16 NIH Institutes and Centers that support the NIH Blueprint for Neuroscience Research; and by the McDonnell Center for Systems Neuroscience at Washington University.

## Author Contributions

S.S. contributed to the problem formulation, design and implementation of the methods, data pre-processing, data analysis and interpretation, and writing the initial draft of the manuscript. D.L. provided the analysis and interpretation of SEEG electrophysiological data. G.C.L. and L.M. contributed to the dMRI preprocessing. E.D. designed the dMRI acquisition protocol and acquired the dMRI data. V.B. performed neurosurgeries, contributed to the acquisition of SEEG data, and provided clinical insights. M.Z. contributed to the subject recruitment and pre-scan assessment. K.D. contributed to the subject recruitment, data interpretation, clinical insights, and provided research guidance. J.P. contributed to the problem formulation, research supervision, acquisition and interpretation of the SEEG data, and writing of the manuscript. J.A.M. contributed to the problem formulation, conception of the project, research supervision, diffusion MRI acquisition, and writing of the manuscript. J.A.M. led the project administration and secured the funding. All authors edited the manuscript and provided feedback on the manuscript and have approved the final version for submission.

## Additional information

**Supplementary information** Supplementary material available in the Extended Data Section A.

**Correspondence and requests for materials** should be addressed to Jennifer A. McNab. For further inquiries regarding this study, please contact the corresponding author.

## References

[1] Park, H.-J., Friston, K.: Structural and functional brain networks: from connections to cognition. Science 342(6158), 1238411 (2013)

[2] Suárez, L.E., Markello, R.D., Betzel, R.F., Misic, B.: Linking structure and function in macroscale brain networks. Trends in Cognitive Sciences 24(4), 302–315 (2020)

[3] Vázquez-Rodríguez, B., Suárez, L.E., Markello, R.D., Shafiei, G., Paquola, C., Hagmann, P., Van Den Heuvel, M.P., Bernhardt, B.C., Spreng, R.N., Misic, B.: Gradients of structure–function tethering across neocortex. Proceedings of the National Academy of Sciences 116(42), 21219–21227 (2019)

[4] Fotiadis, P., Parkes, L., Davis, K.A., Satterthwaite, T.D., Shinohara, R.T., Bassett, D.S.: Structure–function coupling in macroscale human brain networks. Nature Reviews Neuroscience 25(10), 688–704 (2024)

[5] Basser, P.J., Mattiello, J., LeBihan, D.: MR diffusion tensor spectroscopy and imaging. Biophysical Journal 66(1), 259–267 (1994)

[6] Seguin, C., Sporns, O., Zalesky, A.: Brain network communication: concepts, models and applications. Nature Reviews Neuroscience 24(9), 557–574 (2023)

[7] Hamandi, K., Powell, H.R., Laufs, H., Symms, M.R., Barker, G.J., Parker, G.J., Lemieux, L., Duncan, J.: Combined eeg-fmri and tractography to visualise propagation of epileptic activity. Journal of Neurology, Neurosurgery & Psychiatry 79(5), 594–597 (2008)

[8] Shine, J.M., Lewis, L.D., Garrett, D.D., Hwang, K.: The impact of the human thalamus on brain-wide information processing. Nature Reviews Neuroscience 24(7), 416–430 (2023)

[9] Wu, T.Q., Kaboodvand, N., McGinn, R.J., Veit, M., Davey, Z., Datta, A., Graber, K.D., Meador, K.J., Fisher, R., Buch, V., et al.: Multisite thalamic recordings to characterize seizure propagation in the human brain. Brain 146(7), 2792–2802 (2023)

[10] French, J.A.: Refractory epilepsy: clinical overview. Epilepsia 48, 3–7 (2007)

[11] Cardinale, F., Cossu, M., Castana, L., Casaceli, G., Schiariti, M.P., Miserocchi, A., Fuschillo, D., Moscato, A., Caborni, C., Arnulfo, G., et al.: Stereoelectroencephalography: surgical methodology, safety, and stereotactic application accuracy in 500 procedures. Neurosurgery 72(3), 353–366 (2013)

[12] Tomlinson, S.B., Buch, V.P., Armstrong, D., Kennedy, B.C.: Stereoelectroencephalography in pediatric epilepsy surgery. Journal of Korean Neurosurgical Society 62(3), 302–312 (2019)

[13] Buch, V.P., Parvizi, J.: Evolution of seeg strategy: Stanford experience. Neurosurgery Clinics of North America 35(1), 83–85 (2023)

[14] Siddiqi, S.H., Kording, K.P., Parvizi, J., Fox, M.D.: Causal mapping of human brain function. Nature Reviews Neuroscience 23(6), 361–375 (2022)

[15] Matsumoto, R., Nair, D.R., LaPresto, E., Bingaman, W., Shibasaki, H., Lüders, H.O.: Functional connectivity in human cortical motor system: a cortico-cortical evoked potential study. Brain 130(1), 181–197 (2007)

[16] Lyu, D., Stiger, J.R., Lusk, Z., Buch, V., Parvizi, J.: Mapping human thalamocortical connectivity with electrical stimulation and recording. Nature Neuroscience, 1–13 (2025)

[17] Matsumoto, R., Nair, D.R., LaPresto, E., Najm, I., Bingaman, W., Shibasaki, H., Lüders, H.O.: Functional connectivity in the human language system: a cortico-cortical evoked potential study. Brain 127(10), 2316–2330 (2004)

[18] Keller, C.J., Honey, C.J., Megevand, P., Entz, L., Ulbert, I., Mehta, A.D.: Mapping human brain networks with cortico-cortical evoked potentials. Philosophical Transactions of the Royal Society B: Biological Sciences 369(1653) (2014)

[19] Wu, S., Issa, N.P., Rose, S.L., Haider, H.A., Nordli Jr, D.R., Towle, V.L., Warnke, P.C., Tao, J.X.: Depth versus surface: A critical review of subdural and depth electrodes in intracranial electroencephalographic studies. Epilepsia 65(7), 1868–1878 (2024)

[20] Maier-Hein, K.H., Neher, P.F., Houde, J.-C., Cote, M.-A., Garyfallidis, E., Zhong, J., Chamberland, M., Yeh, F.-C., Lin, Y.-C., Ji, Q., et al.: The challenge of mapping the human connectome based on diffusion tractography. Nature communications 8(1), 1349 (2017)

[21] Shailja, S., Chen, J.W., Grafton, S.T., Manjunath, B.: Retrace: Topological evaluation of white matter tractography algorithms using reeb graphs. In: International Workshop on Computational Diffusion MRI, pp. 177–191 (2023). Springer

[22] Yendiki, A., Aggarwal, M., Axer, M., Howard, A.F., Walsum, A.-M.v.C., Haber, S.N.: Post mortem mapping of connectional anatomy for the validation of diffusion mri. NeuroImage 256, 119146 (2022)

[23] Calabrese, E., Badea, A., Cofer, G., Qi, Y., Johnson, G.A.: A diffusion mri tractography connectome of the mouse brain and comparison with neuronal tracer data. Cerebral Cortex 25(11), 4628–4637 (2015)

[24] Silverstein, B.H., Asano, E., Sugiura, A., Sonoda, M., Lee, M.-H., Jeong, J.-W.: Dynamic tractography: Integrating cortico-cortical evoked potentials and diffusion imaging. NeuroImage 215, 116763 (2020)

[25] Crocker, B., Ostrowski, L., Williams, Z.M., Dougherty, D.D., Eskandar, E.N., Widge, A.S., Chu, C.J., Cash, S.S., Paulk, A.C.: Local and distant responses to single pulse electrical stimulation reflect different forms of connectivity. NeuroImage 237, 118094 (2021)

[26] Trebaul, L., Deman, P., Tuyisenge, V., Jedynak, M., Hugues, E., Rudrauf, D., Bhattacharjee, M., Tadel, F., Chanteloup-Foret, B., Saubat, C., et al.: Probabilistic functional tractography of the human cortex revisited. NeuroImage 181, 414–429 (2018)

[27] Seguin, C., Jedynak, M., David, O., Mansour, S., Sporns, O., Zalesky, A.: Communication dynamics in the human connectome shape the cortex-wide propagation of direct electrical stimulation. Neuron 111(9), 1391–1401 (2023)

[28] Abraham, A., Pedregosa, F., Eickenberg, M., Gervais, P., Mueller, A., Kossaifi, J., Gramfort, A., Thirion, B., Varoquaux, G.: Machine learning for neuroimaging with scikit-learn. Frontiers in Neuroinformatics 8, 14 (2014) 10.3389/fninf.2014.00014

[29] Yeh, F.-C.: Population-based tract-to-region connectome of the human brain and its hierarchical topology. Nature Communications 13(1), 4933 (2022)

[30] Aliev, R.A.: Fundamentals of the fuzzy logic-based generalized theory of decisions. Springer (2013)

[31] Nguyen, H.T., Mukaidono, M., Kreinovich, V.: Probability of implication, logical version of bayes theorem, and fuzzy logic operations. In: 2002 IEEE World Congress on Computational Intelligence. 2002 IEEE International Conference on Fuzzy Systems. FUZZ-IEEE’02. Proceedings (Cat. No. 02CH37291), vol. 1, pp. 530–535 (2002). IEEE

[32] Rushton, W.: A theory of the effects of fibre size in medullated nerve. The Journal of Physiology 115(1), 101 (1951)

[33] Drakesmith, M., Harms, R., Rudrapatna, S.U., Parker, G.D., Evans, C.J., Jones, D.K.: Estimating axon conduction velocity in vivo from microstructural mri. NeuroImage 203, 116186 (2019)

[34] Horowitz, A., Barazany, D., Tavor, I., Bernstein, M., Yovel, G., Assaf, Y.: In vivo correlation between axon diameter and conduction velocity in the human brain. Brain Structure and Function 220(3), 1777–1788 (2015)

[35] Chau Loo Kung, G., Knowles, J.K., Batra, A., Ni, L., Rosenberg, J., McNab, J.A.: Quantitative mri reveals widespread, network-specific myelination change during generalized epilepsy progression. NeuroImage 280, 120312 (2023) 10.1016/j.neuroimage.2023.120312

[36] Chau Loo Kung, G., Weber, E.M., Batra, A., Ni, L., Zeineh, M., Chaudhari, A., Adeli, E., Knowles, J.K., McNab, J.A.: Non-parametric prediction of brain mri microstructure using transfer learning. Imaging Neuroscience 3, 00548 (2025)

[37] Zhang, H., Schneider, T., Wheeler-Kingshott, C.A., Alexander, D.C.: Noddi: practical in vivo neurite orientation dispersion and density imaging of the human brain. NeuroImage 61(4), 1000–1016 (2012)

[38] Togo, M., Lyu, D., Huang, W., Pantis, S., Fisher, R., Matsumoto, R., Buch, V., Parvizi, J.: Electrophysiological connections linking medial pulvinar, anterior nuclei of the thalamus and the hippocampus. Brain 148(12), 4315–4324 (2025)

[39] Azeem, A., Ellenrieder, N., Royer, J., Frauscher, B., Bernhardt, B., Gotman, J.: Integration of white matter architecture to stereo-eeg better describes epileptic spike propagation. Clinical Neurophysiology 146, 135–146 (2023)

[40] Giampiccolo, D., Dijk, J., Granados, A., Xiao, F., Fiore, G., Rodionov, R., Li, K., Lysomirski, A.L., McEvoy, A.W., Diehl, B., et al.: Mapping white matter tracts with seeg electrodes. Epilepsia (2025)

[41] David, O., Job, A.-S., De Palma, L., Hoffmann, D., Minotti, L., Kahane, P.: Probabilistic functional tractography of the human cortex. NeuroImage 80, 307–317 (2013)

[42] Stieger, J.R., Pinheiro-Chagas, P., Fang, Y., Li, J., Lusk, Z., Perry, C.M., Girn, M., Contreras, D., Chen, Q., Huguenard, J.R., et al.: Cross-regional coordination of activity in the human brain during autobiographical self-referential processing. Proceedings of the National Academy of Sciences 121(32), 2316021121 (2024)

[43] Middlebrooks, E.H., Grewal, S.S., Stead, M., Lundstrom, B.N., Worrell, G.A., Van Gompel, J.J.: Differences in functional connectivity profiles as a predictor of response to anterior thalamic nucleus deep brain stimulation for epilepsy: a hypothesis for the mechanism of action and a potential biomarker for outcomes. Neurosurgical Focus 45(2), 7 (2018)

[44] Bell, P.T., Shine, J.M.: Subcortical contributions to large-scale network communication. Neuroscience & Biobehavioral Reviews 71, 313–322 (2016)

[45] Aggleton, J.P., O’Mara, S.M.: The anterior thalamic nuclei: core components of a tripartite episodic memory system. Nature Reviews Neuroscience 23(8), 505–516 (2022)

[46] Beaulieu, C.: The basis of anisotropic water diffusion in the nervous system–a technical review. NMR in Biomedicine: An International Journal Devoted to the Development and Application of Magnetic Resonance In Vivo 15(7-8), 435–455 (2002)

[47] Abe, S., Vidmark, J., Hernandez-Martin, E., Kasiri, M., Sorouhmojdehi, R., Seyyed Mousavi, S.A., Sanger, T.D.: Diffusion tractography predicts deep brain stimulation evoked potential amplitude and delay. medRxiv, 2024–04 (2024)

[48] Jbabdi, S., Sotiropoulos, S.N., Savio, A.M., Graña, M., Behrens, T.E.: Model-based analysis of multishell diffusion mr data for tractography: How to get over fitting problems. Magnetic Resonance in Medicine 68(6), 1846–1855 (2012)

[49] Behrens, T.E., Berg, H.J., Jbabdi, S., Rushworth, M.F., Woolrich, M.W.: Probabilistic diffusion tractography with multiple fibre orientations: What can we gain? NeuroImage 34(1), 144–155 (2007)

[50] Parvizi, J., Kastner, S.: Human intracranial eeg: promises and limitations. Nature Neuroscience 21(4), 474 (2018)

[51] Guo, Z.-h., Zhao, B.-t., Toprani, S., Hu, W.-h., Zhang, C., Wang, X., Sang, L., Ma, Y.-s., Shao, X.-q., Razavi, B., et al.: Epileptogenic network of focal epilepsies mapped with cortico-cortical evoked potentials. Clinical Neurophysiology 131(11), 2657–2666 (2020)

[52] Lusk, Z., Kwon, A.Y., Pantis, S., Nam, S., Lyu, D., Fisher, R., Staalduinen, E.K., Buch, V., Parvizi, J.: Combining clinical evaluations and neuroscience research in the human intracranial electroencephalography practice: 15-year cohort study. Journal of Cognitive Neuroscience 37(11), 2108–2125 (2025)

[53] Groppe, D.M., Bickel, S., Dykstra, A.R., Wang, X., Mégevand, P., Mercier, M.R., Lado, F.A., Mehta, A.D., Honey, C.J.: ielvis: An open source matlab toolbox for localizing and visualizing human intracranial electrode data. Journal of Neuroscience Methods 281, 40–48 (2017)

[54] Fischl, B.: Freesurfer. NeuroImage 62(2), 774–781 (2012)

[55] Papademetris, X., Jackowski, M.P., Rajeevan, N., DiStasio, M., Okuda, H., Constable, R.T., Staib, L.H.: Bioimage suite: An integrated medical image analysis suite: An update. The Insight Journal 2006, 209 (2006)

[56] McInnes, L., Healy, J., Melville, J.: UMAP: Uniform manifold approximation and projection for dimension reduction. arXiv preprint arXiv:1802.03426 (2018)

[57] Daducci, A., Canales-Rodríguez, E.J., Zhang, H., Dyrby, T.B., Alexander, D.C., Thiran, J.-P.: Accelerated microstructure imaging via convex optimization (amico) from diffusion mri data. NeuroImage 105, 32–44 (2015)

[58] Kumar, V.J., Van Oort, E., Scheffler, K., Beckmann, C.F., Grodd, W.: Functional anatomy of the human thalamus at rest. NeuroImage 147, 678–691 (2017)

