## Extended Data for "Diffusion MRI Tractography Predicts Electrophysiological Connectivity and Explains Spectral Signatures of Evoked Potentials in the Human Brain"

### Appendix A Extended Data

**Table A1**: Participant Demographics

| Subject ID | Gender | Age | # Electrode contacts | Hemisphere coverage | # $E$ Pairs | # $\neg E$ Pairs |
|---|---|---|---|---|---|---|
| S01_166* | F | 38 | 176 | B | 1425 | 664 |
| S02_167* | F | 23 | 204 | R | 2677 | 903 |
| S03_169* | M | 23 | 186 | B | 2591 | 1613 |
| S04_170 | M | 41 | 116 | R | 1231 | 215 |
| S05_171 | M | 52 | 252 | B | 4035 | 1393 |
| S06_172* | M | 47 | 174 | B | 2644 | 1373 |
| S07_176* | F | 34 | 156 | L | 2138 | 766 |
| S08_177* | M | 19 | 168 | B | 1942 | 893 |
| S09_178* | M | 28 | 252 | B | 3706 | 909 |
| S10_181* | F | 23 | 310 | B | 918 | 213 |
| S11_182 | F | 38 | 260 | B | 3391 | 1059 |
| S12_183* | M | 33 | 174 | B | 2117 | 1169 |
| S13_185 | F | 36 | 142 | B | 2357 | 853 |
| S14_188 | M | 31 | 122 | B | 2441 | 2249 |
| S15_189 | M | 27 | 184 | B | 1031 | 61 |
| S16_190 | F | 50 | 116 | R | 1180 | 684 |
| S17_192 | M | 20 | 88 | L | 128 | 15 |
| S18_193 | M | 40 | 184 | B | 3138 | 5092 |
| S19_194 | M | 38 | 246 | B | 4917 | 733 |
| S20_195 | F | 47 | 138 | B | 1620 | 678 |
| S21_197 | M | 20 | 214 | B | 4944 | 2137 |
| S22_198 | M | 39 | 204 | B | 4259 | 1009 |
| S23_199 | M | 37 | 178 | B | 4015 | 1024 |
| S24_201 | M | 28 | 188 | B | 2606 | 1783 |
| S25_202 | F | 49 | 194 | B | 3253 | 1306 |
| S26_205 | F | 49 | 184 | B | 4663 | 1053 |
| S27_206 | M | 59 | 122 | B | 1280 | 489 |
| S28_207 | F | 67 | 112 | L | 1638 | 471 |
| S29_208 | M | 58 | 250 | B | 8991 | 1857 |
| S30_209 | M | 47 | 248 | B | 4857 | 1280 |
| S31_210* | M | 52 | 204 | B | 3294 | 722 |
| S32_211 | F | 32 | 198 | B | 3971 | 1288 |
| S33_212 | M | 36 | 218 | B | 2084 | 357 |
| S34_213 | M | 20 | 216 | B | 4922 | 1391 |
| S35_214 | F | 28 | 184 | B | 1617 | 206 |
| S36_218 | F | 56 | 206 | B | 718 | 13 |
| S37_220 | M | 32 | 216 | B | 4431 | 1469 |
| S38_221 | F | 44 | 234 | B | 3556 | 681 |
| S39_222 | F | 49 | 176 | B | 1160 | 151 |
| S40_236 | M | 26 | 218 | B | 3244 | 709 |

[1]* indicates subjects with mapped diffusion MRI data. B: bilateral; L: left; R: right. # $E$ pairs: number of electrode pairs labeled as electrophysiologically connected. # $\neg E$ pairs: number of electrode pairs labeled as electrophysiologically not connected.

Table (A2) HCP Subjects Demographics

| Subject ID | Age | Gender |
|---|---|---|
| 100206 | 26-30 | M |
| 100307 | 26-30 | F |
| 100408 | 31-35 | M |
| 100610 | 26-30 | M |
| 101006 | 31-35 | F |
| 101107 | 22-25 | M |
| 101309 | 26-30 | M |
| 101410 | 26-30 | M |
| 101915 | 31-35 | F |
| 102008 | 22-25 | M |
| 102109 | 26-30 | M |
| 102311 | 26-30 | F |
| 102513 | 26-30 | M |
| 102614 | 22-25 | M |
| 102715 | 26-30 | M |
| 102816 | 26-30 | F |
| 103010 | 22-25 | M |
| 103111 | 26-30 | M |
| 103212 | 31-35 | M |
| 103414 | 22-25 | F |
| 103515 | 26-30 | F |
| 103818 | 31-35 | F |
| 104012 | 26-30 | F |
| 104416 | 31-35 | F |
| 104820 | 36+ | F |
| 105014 | 26-30 | F |
| 105115 | 31-35 | M |
| 105216 | 26-30 | M |
| 105620 | 31-35 | F |
| 105923 | 31-35 | F |
| 106016 | 31-35 | F |
| 106319 | 26-30 | M |
| 106521 | 26-30 | F |
| 106824 | 22-25 | M |
| 107018 | 26-30 | F |
| 107321 | 22-25 | F |
| 107422 | 22-25 | M |
| 107725 | 31-35 | F |
| 108020 | 22-25 | M |
| 108121 | 26-30 | F |
| 108222 | 31-35 | M |
| 108323 | 26-30 | F |
| 108525 | 22-25 | M |
| 108828 | 31-35 | M |
| 109123 | 31-35 | M |
| 109830 | 31-35 | F |
| 110007 | 31-35 | F |
| 110411 | 31-35 | M |
| 110613 | 26-30 | M |
| 111009 | 26-30 | F |
| 111211 | 26-30 | F |
| 111312 | 31-35 | F |
| 111413 | 26-30 | F |
| 111514 | 31-35 | M |
| 111716 | 31-35 | F |
| 112112 | 26-30 | M |
| 112314 | 26-30 | F |
| 112516 | 31-35 | F |
| 112819 | 26-30 | F |
| 112920 | 31-35 | M |
| 113215 | 26-30 | F |
| 113316 | 22-25 | M |
| 113619 | 31-35 | F |
| 113821 | 26-30 | F |
| 113922 | 31-35 | M |
| 114116 | 26-30 | F |
| 114217 | 26-30 | F |
| 114318 | 22-25 | F |
| 114419 | 31-35 | M |
| 114621 | 26-30 | M |
| 114823 | 31-35 | F |
| 115017 | 31-35 | F |
| 115219 | 31-35 | M |
| 115320 | 31-35 | F |
| 115724 | 22-25 | F |
| 115825 | 22-25 | M |
| 116221 | 22-25 | M |
| 116423 | 26-30 | M |
| 116524 | 26-30 | M |
| 116726 | 26-30 | M |
| 117021 | 26-30 | F |
| 117122 | 26-30 | F |
| 117324 | 22-25 | F |
| 117930 | 31-35 | F |
| 118023 | 26-30 | F |
| 118124 | 31-35 | F |
| 118225 | 26-30 | M |
| 118528 | 26-30 | F |
| 118730 | 22-25 | M |
| 118831 | 22-25 | M |
| 118932 | 26-30 | M |
| 119025 | 26-30 | M |
| 119126 | 22-25 | F |
| 119732 | 31-35 | F |
| 119833 | 26-30 | F |
| 120010 | 26-30 | F |
| 120111 | 26-30 | F |
| 120212 | 31-35 | F |
| 120414 | 26-30 | F |
| 120515 | 26-30 | F |
| 120717 | 31-35 | F |
| 121416 | 26-30 | M |
| 121618 | 31-35 | M |
| 121719 | 31-35 | F |
| 121921 | 31-35 | M |
| 122317 | 31-35 | M |
| 122418 | 26-30 | F |
| 122620 | 26-30 | M |
| 122822 | 31-35 | F |
| 123117 | 26-30 | M |
| 123420 | 26-30 | F |
| 123521 | 22-25 | M |
| 123723 | 31-35 | F |
| 123824 | 22-25 | M |
| 123925 | 26-30 | F |
| 124220 | 31-35 | M |
| 124422 | 31-35 | F |
| 124624 | 26-30 | F |
| 124826 | 31-35 | F |
| 125222 | 26-30 | M |
| 125424 | 31-35 | F |
| 125525 | 31-35 | F |
| 126325 | 26-30 | F |
| 126426 | 31-35 | F |
| 126628 | 31-35 | M |
| 127226 | 22-25 | M |
| 127327 | 22-25 | M |
| 127630 | 22-25 | F |
| 127731 | 31-35 | F |
| 127832 | 31-35 | F |
| 127933 | 31-35 | M |
| 128026 | 26-30 | F |
| 128127 | 26-30 | F |
| 128632 | 31-35 | F |
| 128935 | 31-35 | M |
| 129028 | 26-30 | M |
| 129129 | 31-35 | F |
| 129331 | 26-30 | F |
| 129634 | 22-25 | M |
| 129937 | 26-30 | F |
| 130013 | 26-30 | M |
| 130114 | 26-30 | M |
| 130316 | 26-30 | F |
| 130417 | 26-30 | M |
| 130518 | 31-35 | F |
| 130619 | 26-30 | F |
| 130720 | 31-35 | M |
| 130821 | 22-25 | F |
| 130922 | 31-35 | F |
| 131217 | 26-30 | F |
| 131419 | 26-30 | F |
| 131722 | 26-30 | F |
| 131823 | 26-30 | M |
| 131924 | 22-25 | M |
| 132017 | 22-25 | F |
| 132118 | 26-30 | F |
| 133019 | 26-30 | F |
| 133625 | 26-30 | M |
| 133827 | 22-25 | M |
| 133928 | 26-30 | M |
| 134021 | 31-35 | F |
| 134223 | 26-30 | F |
| 134324 | 26-30 | M |
| 134425 | 26-30 | F |
| 134627 | 26-30 | M |
| 134728 | 26-30 | M |
| 134829 | 31-35 | F |
| 135124 | 31-35 | F |
| 135225 | 22-25 | F |
| 135528 | 26-30 | F |
| 135629 | 22-25 | M |
| 135730 | 22-25 | M |
| 135932 | 26-30 | F |
| 136126 | 26-30 | M |
| 136227 | 31-35 | M |
| 136631 | 22-25 | M |
| 136732 | 31-35 | F |
| 136833 | 31-35 | M |
| 137027 | 31-35 | M |
| 137128 | 31-35 | F |
| 137229 | 22-25 | M |
| 137431 | 31-35 | F |
| 137532 | 31-35 | M |
| 137633 | 22-25 | F |
| 137936 | 31-35 | M |
| 138130 | 31-35 | F |
| 138231 | 31-35 | F |
| 138332 | 26-30 | M |
| 138534 | 22-25 | M |
| 138837 | 22-25 | M |
| 139233 | 31-35 | F |
| 139435 | 26-30 | F |
| 139637 | 31-35 | F |
| 139839 | 26-30 | M |
| 140117 | 26-30 | F |
| 140319 | 36+ | F |
| 140420 | 26-30 | F |
| 140824 | 31-35 | M |
| 140925 | 22-25 | F |
| 141119 | 26-30 | M |

**Table A3**: $P(S \mid E)$ for "direct" connectivity across increasing $F1$ thresholds $th_{r^*}$, shown for $s_1$-connectivity for $th_{s_1} = 0$ and $s_2$-connectivity definitions for $th_{s_2} = 0$ and non-zero overlap with the tract region in HCP atlas.

| $th_{r^*}$ for F1 | $P(S\|E)$ for $s_1$-connectivity | $P(S\|E)$ for $s_2$-connectivity |
|---|---|---|
| 0.1 | 0.38 | 0.67 |
| 0.2 | 0.39 | 0.69 |
| 0.3 | 0.42 | 0.71 |
| 0.4 | 0.45 | 0.74 |
| 0.5 | 0.5 | 0.77 |
| 0.6 | 0.57 | 0.82 |
| 0.7 | 0.68 | 0.89 |
| 0.8 | 0.78 | 0.86 |

**Table A4**: $s_1$ mixed-effects model results for direct versus indirect pathway comparisons. Coefficients are indirect minus direct estimates from models of the form `value ~ is_indirect + (1 | subject_id)`.

| Comparison | Feature | Subjects | Observations | Coef. | 95% CI | $p$ | BH $p$ |
|---|---|---|---|---|---|---|---|
| C→C vs C→C→C | F1 | 9 | 3,994 | -0.180 | [-0.204, -0.156] | 2.84e-47 | 3.41e-46 |
| C→C vs C→C→C | F2 | 9 | 3,873 | -0.039 | [-0.059, -0.020] | 8.36e-05 | 2.51e-04 |
| C→C vs C→C→C | F3 | 9 | 3,778 | 0.038 | [0.014, 0.061] | 0.00162 | 0.00323 |
| C→C vs C→T→C | F1 | 6 | 2,806 | -0.307 | [-0.388, -0.226] | 9.01e-14 | 5.40e-13 |
| C→C vs C→T→C | F2 | 8 | 3,228 | -0.060 | [-0.122, 0.003] | 0.0627 | 0.0836 |
| C→C vs C→T→C | F3 | 8 | 3,105 | 0.117 | [0.045, 0.190] | 0.00141 | 0.00323 |
| T→C vs T→C→C | F1 | 10 | 786 | -0.133 | [-0.174, -0.093] | 9.90e-11 | 3.96e-10 |
| T→C vs T→C→C | F2 | 10 | 702 | -0.035 | [-0.072, 0.001] | 0.0588 | 0.0836 |
| T→C vs T→C→C | F3 | 10 | 798 | -0.019 | [-0.069, 0.030] | 0.450 | 0.450 |
| T→T vs T→C→T | F1 | 3 | 57 | -0.147 | [-0.315, 0.020] | 0.0845 | 0.101 |
| T→T vs T→C→T | F2 | 3 | 57 | -0.117 | [-0.265, 0.031] | 0.123 | 0.134 |
| T→T vs T→C→T | F3 | 3 | 59 | -0.185 | [-0.339, -0.031] | 0.0182 | 0.0312 |

**Table A5**: Per-subject Pearson correlation of conduction velocity with the diffusion microstructure properties: FA/MD/NDI/FWF/ODI

| Subject ID | FA | MD | NDI | FWF | ODI | #pairs |
|---|---|---|---|---|---|---|
| S01_166 | 0.45 | -0.39 | 0.40 | 0.49 | -0.45 | 234 |
| S02_167 | 0.45 | -0.37 | 0.37 | 0.23 | -0.53 | 374 |
| S03_169 | 0.33 | -0.31 | 0.29 | 0.09 | -0.27 | 195 |
| S06_172 | 0.50 | -0.31 | 0.27 | 0.04 | -0.46 | 324 |
| S07_176 | -0.05 | 0.07 | -0.10 | -0.11 | -0.04 | 347 |
| S08_177 | 0.39 | -0.20 | 0.17 | 0.05 | -0.41 | 349 |
| S09_178 | 0.50 | -0.35 | 0.37 | 0.30 | -0.49 | 512 |
| S10_181 | 0.14 | -0.01 | 0.00 | -0.04 | -0.19 | 101 |
| S12_183 | 0.24 | -0.07 | 0.10 | 0.14 | -0.30 | 237 |
| S31_210 | 0.37 | -0.29 | 0.30 | 0.22 | -0.38 | 353 |

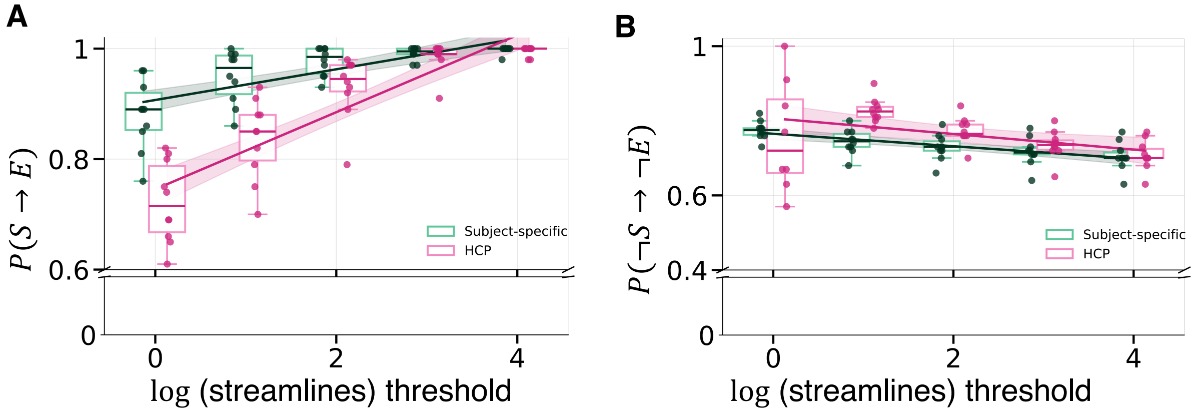


**Fig. A1**: **Structural connectivity derived from patient-normative dMRI from 200 HCP subjects predicts the causal electrophysiological connectivity.** **(A)** Probability of electrophysiological connectivity given structural connectivity, $P(S \to E)$, as a function of the $\log_{10}$(streamline) threshold. **(B)** Probability of the absence of electrophysiological connectivity given the absence of structural connectivity, $P(\neg S \to \neg E)$, as a function of the $\log_{10}$(streamline) threshold. Structural connectivity was evaluated using subject-specific diffusion MRI tractography (green) and the HCP atlas (pink). Dots represent individual participants. Solid lines show linear fits across participants and shaded regions indicate 95% confidence intervals.

**Table A6**: Patient-specific structural connectivity versus Non-patient-specific structural connectivity implying causal electrophysiological connectivity ($P(S \to E)$) across streamline count thresholds.

| $\text{th}_{s_1}$ | Patient-specific | | | Non-specific | | |
|---|---|---|---|---|---|---|
| | $\#(S \cap E)$ | $\#(S \cap \neg E)$ | $P(S \to E)$ | $\#(S \cap E)$ | $\#(S \cap \neg E)$ | $P(S \to E)$ |
| 0 | 786 | 100 | 0.89 | 674 | 84 | 0.89 |
| 10 | 407 | 24 | 0.94 | 344 | 20 | 0.95 |
| 100 | 253 | 6 | 0.98 | 211 | 2 | 0.99 |
| 1000 | 146 | 1 | 0.99 | 126 | 0 | 1.00 |
| 10000 | 60 | 0 | 1.00 | 44 | 0 | 1.00 |

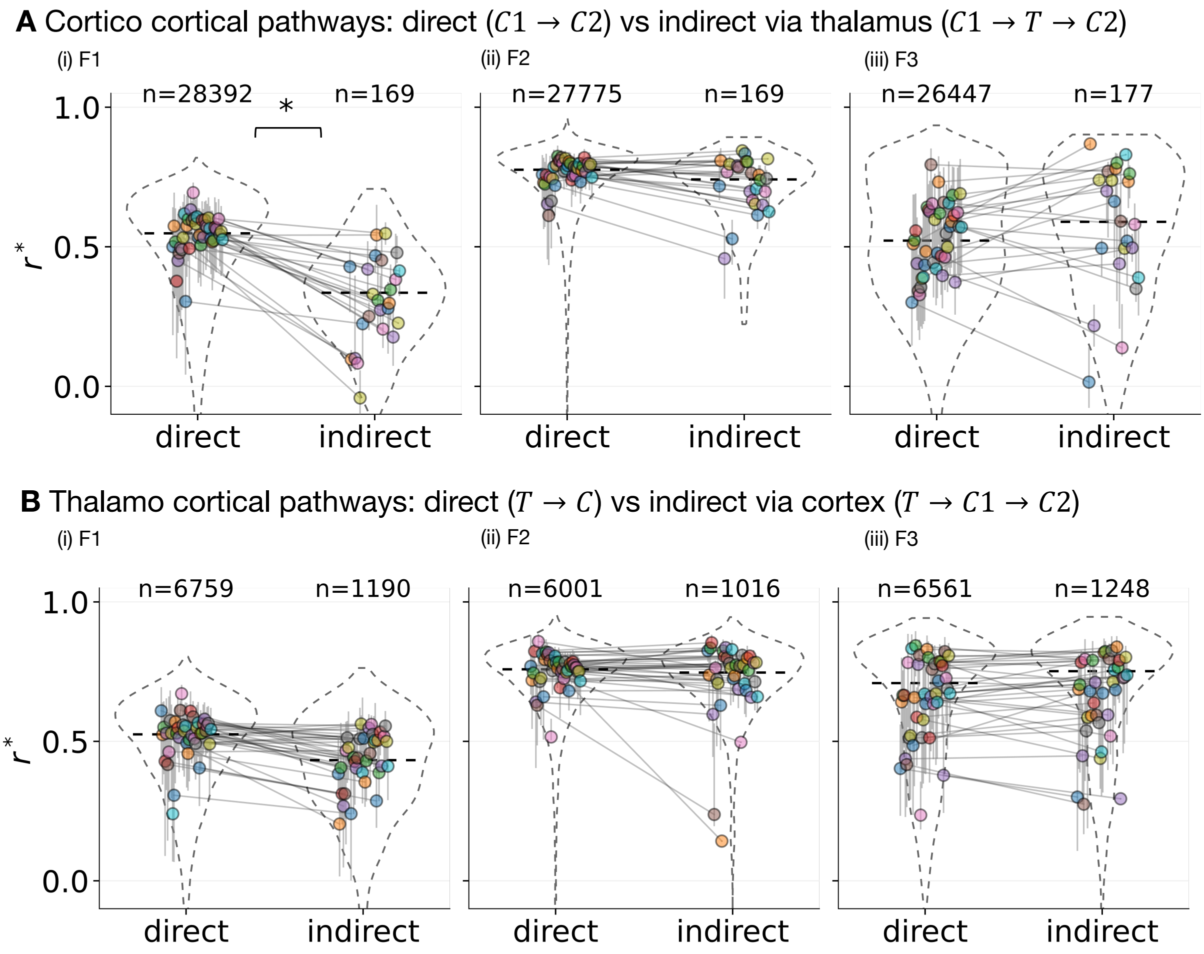


**Fig. A2**: **Comparison of spectral features of the evoked potentials in direct and indirect structural pathways under the $s_2$-connectivity definition ($N$ = 40 subjects). (A)** Corticocortical pathways with indirect connections via thalamus: (i) Correlation with F1 spectral features significantly reduce in indirect pathways relative to direct pathways. (ii, iii) F2 and F3 spectral features do not differ significantly between direct and indirect pathways. **(B)** Thalamocortical pathways with indirect connections via cortex: (i) Correlations with F1 spectral features reduce in indirect pathways relative to direct pathways but not significantly. (ii, iii) F2 and F3 spectral features do not differ significantly between direct and indirect pathways. Statistical significance between direct vs. indirect connections is indicated by asterisks (* $p < 0.05$, absolute difference in mean $> 0.1$). Each colored circle is the subject-wise median of peak correlations and vertical bars are the inter-quartile range.
